# The impact of strain heterogeneity on genomic profiling accuracy in mixed-strain *Escherichia coli* metagenomes

**DOI:** 10.64898/2026.10.19.760091

**Authors:** Vaishnavi Divya Shridar, Stefan Engblom

**Author notes:** Corresponding author.* URL: https://stefanengblom.github.io.

## Abstract

Metagenomic sequencing is increasingly used for surveillance of bacterial populations, enabling simultaneous characterisation of antimicrobial resistance, virulence, and epidemiologically relevant strain markers. However, the ability of commonly used bioinformatic tools to recover these features from heterogeneous populations remains incompletely understood.

Here, we developed a controlled benchmarking framework based on simulated single-species *Escherichia coli* metagenomes generated from genomes with PCR-determined annotations. Assembly-based and read-based tools were evaluated for antimicrobial resistance gene (ARG) detection, virulence gene (VG) detection, multilocus sequence typing (MLST), and serotype prediction. In addition, datasets with varying strain compositions were used to assess the effect of population complexity on strain-level characterisation.

We found that recovery of individual genetic determinants of interest (7 ARGs, 3 VGs) was consistently more successful than recovery of strain-level markers. ARG detection achieved high recovery rates across all evaluated tools (84.3–90.0%), while short read sequencing typer SRST2 achieved the highest virulence gene recovery rate (100.0%). In contrast, MLST and serotype prediction were strongly affected by strain composition. None of the evaluated MLST tools correctly assigned the dominant sequence type in the multi-strain log-normal dataset, despite successful partial recovery of MLST alleles, while complete serotype recovery remained uncommon across all methods. Decreasing the complexity of the simulated metagenome by increasing the abundance of the dominant strain improved both MLST and serotype prediction performance, while long-read analyses indicated that increased read length was insufficient to overcome the challenges associated with mixed-strain communities.

These findings demonstrate that performance estimates derived from gene detection do not necessarily translate to strain-level characterisation. While individual resistance and virulence determinants could often be recovered reliably, accurate reconstruction of sequence types and serotypes remained strongly constrained by strain heterogeneity. More broadly, the results highlight current limitations in resolving strain composition from metagenomic data and provide a framework for investigating the abundance thresholds and population structures required for reliable strain-resolved surveillance.

**Contribution Statement:** This study quantifies how strain heterogeneity influences recovery of clinically relevant genomic features from low-complexity *Escherichia coli* metagenomes. We show that gene-level characterisation remains largely robust, while strain-level markers such as MLST and serotype are substantially more sensitive to mixed-strain populations. These findings highlight strain heterogeneity as a major barrier to linking clinically relevant genes to their epidemiological background in metagenomic surveillance.

## 1 Background

In the field of pathogen surveillance, wastewater-based epidemiology (WBE) has emerged as a cost-effective and scalable tool (1). It has been demonstrated that such population-level surveillance has allowed for viral RNA to be detected before the later reporting of clinical cases (2). Given the success of WBE in viral surveillance, it is now of great interest to apply this tool to bacterial targets (3). This urgency is exacerbated by multi-drug resistant bacteria increasing in prevalence and little successful antibiotic development, making antimicrobial resistance (AMR) a major concern in current times (4). Notably, in 2022, *E. coli* was identified as the leading bacterial pathogen for deaths associated with antibiotic resistance (5).

From a surveillance perspective, the accurate identification of genomic characteristics linked to antimicrobial resistance and pathogenicity is essential for the assessment of public health risk. In *E. coli*, four characteristics of pathogenic and antimicrobial resistance potential are commonly identified in practice: (1) antibiotic resistance genes (ARGs), (2) virulence genes (VGs), (3) bacterial multi-locus sequence type (MLST) and (4) serotype (6–9). ARGs encode pathways that confer resistance to antibiotics, while virulence genes (VGs) encode virulence factors (VFs) that contribute to pathogenic mechanisms such as host colonisation, invasion, immune evasion and host damage (10, 11). Although metagenomic sequencing primarily identifies the presence of these genes rather than their expression or activity, ARGs and VGs are widely used as important determinants of bacterial resistance to antibiotics, and disease severity in infected humans, respectively (12). Multilocus sequence types (STs) are discrete genotypic classifications defined by unique combinations of allele sequences at multiple housekeeping loci while serotyping presents a phenotypic classification of microorganisms based on antigenic variation in cell surface structures (13, 14). Combining these complementary typing frameworks improves the resolution when distinguishing epidemiologically related strains from unrelated lineages sharing similar antigenic profiles (15). Together, these four characteristics enable comprehensive characterisation of bacterial populations and their associated risks in infected humans.

Metagenomic sequencing has become a powerful tool for investigating these characteristics directly from environmental samples, including wastewater. Unlike isolate-based sequencing, metagenomics allows simultaneous detection of multiple strains and gene classes within complex microbial communities. However, analysis of genomic signals from multiple organisms presents significant analytical challenges, particularly for strain-level inference and the accurate identification of clinically relevant markers. Recent progress in metagenomic sequencing has spurred greater dependence on bioinformatics tools rather than targeted polymerase chain reaction (PCR) for bacterial typing and gene detection in complex matrices such as wastewater (16). This has been due to falling sequencing costs, improved sequencing accuracy and the large availability of publicly accessible metagenomes (17). Many commonly used bioinformatic tools for MLST, serotyping, and gene detection were originally developed and validated using isolate whole-genome sequencing data, where assumptions of genome completeness and single-strain representation are typically satisfied. Their performance under metagenomic conditions where these assumptions are violated remains insufficiently characterised.

Between isolate sequencing and the more complex metagenomics exists an important and under-explored scenario of low-complexity and single-species metagenomes derived from selective cultivation or colony pooling. In such datasets, a dominant *E. coli* strain may be present, but residual heterogeneity persists due to co-occurring strains. These conditions are highly relevant to wastewater surveillance workflows, where enrichment-based approaches are often used to increase target organism abundance, ensuring commensal signals are not overwhelmed by target genes and that inconclusive results or false negatives are avoided (18). This intermediate system challenges both isolate-focused and metagenome-specific tools, providing a useful framework for evaluating their robustness across input types.

Despite the growing reliance on bioinformatic tools for genomic profiling, systematic benchmarking under controlled conditions remains limited. Benchmarking is recognised as a critical component of method development and validation, but many studies are conducted in a fragmented or task-specific manner (19, 20). While there has been some evaluation of *in silico* tools for direct metagenomes, tool performance remains dependent on dataset characteristics and underlying database composition, thus requiring re-evaluation (21, 22). Existing studies often depend on bioinformatically derived annotations, highlighting the importance of controlled datasets and careful benchmarking design to ensure reliable performance evaluation (23). Few studies have evaluated tool performance across multiple clinically relevant feature classes within a unified framework, as most benchmarking studies focus on specific analytical tasks such as taxonomic classification and demonstrate substantial variability in tool performance across identical datasets (20, 23). This highlights the need for bench-marking approaches that incorporate independent, experimentally derived ground truth and reflect realistic surveillance scenarios in a unified framework.

In this study, we address this gap by developing a controlled benchmarking framework using simulated short-read single-species *E. coli* metagenomes derived from genomes with PCR-determined annotations. We evaluate a panel of widely used tools spanning read-based, assembly-based, and metagenomic approaches for their ability to recover antimicrobial resistance genes, virulence determinants, MLST and serotypes. We hypothesise that tools designed for isolate sequencing can recover dominant strain features under low-complexity conditions, but that their performance will vary according to underlying methodological assumptions of the individual tools. By comparing the two major metagenomic analysis schools of read-based and assembly-based approaches within a controlled benchmarking framework, we aim to assess their robustness under conditions representative of wastewater surveillance (24). In this context, our objective is not only to compare tool accuracy, but also to identify systematic biases and failure modes when methods are applied outside their original design space. This positions our work as a benchmark for assessing the reliability and transferability of genomic profiling tools across different data types encountered in public health microbiology.

## 2 Methods and Materials

To simulate low-complexity metagenomic conditions, relevant input genomes with PCR-annotations were required. Section 2.1 introduces the whole genome selection process and the construction of three different datasets for comprehensive evaluation of gene detection and strain typing. Section 2.2 details the metagenome simulation methodology and the preprocessing steps applied to the simulated metagenomes. Section 2.3 reports the choice and implementation of bioinformatic tools, as well as the performance metrics used for comparing tool predictions against the PCR-derived ground truth. Figure 2.1 presents an overview of the complete benchmarking framework from data selection to performance evaluation.

**Figure 2.1:**
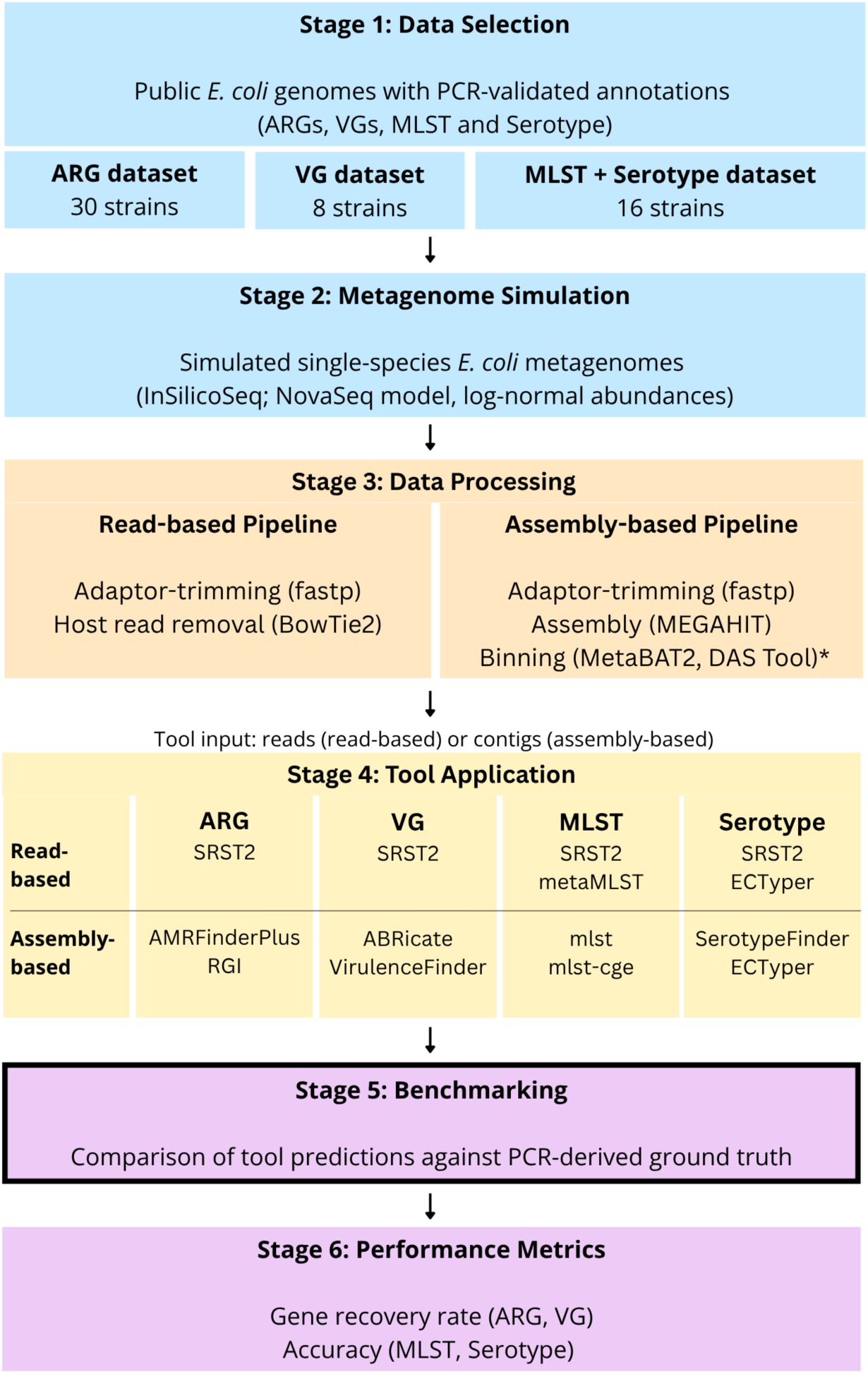
Overview of the benchmarking framework showing dataset construction from *E. coli* genomes with experimental annotations derived from in-lab polymerase chain reactions (PCR), simulation of low-complexity metagenomes, and evaluation of genomic profiling tools across methodological classes using the experimentally derived ground truth. *As binning consistently reduced recovery of clinical characteristics of interest, the assembly-based workflow and results focused on analyses performed directly on assembled contigs.

### 2.1 Clinical Characteristics and Genomes of Interest

Four classes of genomic features relevant to *E. coli* surveillance were evaluated: antimicrobial resistance genes (ARGs), virulence genes, sequence types (STs), and serotypes. Target ARGs included *bla_TEM_*, *sul1*, *sul2*, *sul3*, *tetA*, *tetB* and *tetD* representing commonly observed resistance determinants in *E. coli* across beta-lactam, sulfonamide, and tetracycline classes (25). These genes were selected based on their relevance to clinical and environmental surveillance and the availability of experimentally validated annotations, where gene presence had been confirmed by PCR in the source isolates (26). The selected virulence genes are *stx1*, *stx2*, and *eae*, representing key functional categories of toxin production, host adhesion, and capsule formation (27). These genes are widely used markers of pathogenic potential in *E. coli* and were chosen based on their prevalence in surveillance studies and availability of PCR-confirmed gene annotations in the corresponding reference genomes. Sequence typing was performed using the Achtman 7-gene MLST scheme, and serotyping was based on O- and H-antigen classification (14, 28). Feature selection was guided by the availability of independently validated PCR metadata associated with the source isolates and their relevance to clinical and environmental surveillance.

Publicly available *E. coli* genomes with experimentally validated PCR annotations were selected to ensure independent ground truth. Table 2.1 summarises the composition of the simulated datasets used for benchmarking. Three feature-specific datasets were constructed: (1) ARG dataset from 30 strains capturing diverse resistance determinants, (2) VG dataset from 8 strains with curated virulence gene profiles and (3) MLST+Serotype dataset from 16 strains representing distinct ST-serotype combinations. Dataset sizes were selected to reflect differences in biological diversity and redundancy across feature classes, with larger datasets used for ARG detection due to greater gene variation. A summary of all genomes used is provided in Supplementary Table S1, indicating their corresponding dataset, genome accession number and PCR-validated annotations.

**Table 2.1:** Summary of simulated *E. coli* datasets used for benchmarking genomic profiling tools. For each of the three datasets (ARG, VG and MLST+Serotype), the number of whole genome strains used in the metagenome simulation and the features to be predicted from the dataset is indicated.

| Dataset | Strains (n) | Features | Rationale |
| --- | --- | --- | --- |
| ARG | 30 | *bla*<sub>TEM</sub>, *sul1*, *sul2*,<br>*sul3*, *tetA*, *tetB*, *tetD* | Captures diversity of antimicrobial resistance genes for robust benchmarking. |
| VG | 8 | *stx1*, *stx2*, *eae* | PCR-validated targets for controlled assessment of detection accuracy. |
| MLST + Serotype | 16 | MLST alleles and O/H antigen loci | Representative diversity supports evaluation of strain-typing performance. |

### 2.2 Metagenome Simulation and Processing

Single-species metagenomes were simulated using `InSilicoSeq` (v2.0.1) (29). Paired-end reads (150 bp) were generated using a NovaSeq error model at 2 million reads per paired-end. For each dataset (ARG, VG, MLST+Serotype), 10 metagenomes were simulated, resulting in a total of 30 metagenomic samples. Relative abundances of strains within each metagenome were modelled using a log-normal distribution to reflect realistic variation in strain dominance. Each metagenome consisted exclusively of *E. coli* strains, representing a low-complexity community with one or more dominant strains, consistent with enrichment-based wastewater surveillance workflows.

To evaluate whether the analytical strategy influenced strain characterisation and gene detection performance, both read-based and assembly-based workflows were implemented.

Metagenomic analyses are commonly performed using either assembly-based or assembly-free (read-based) approaches (24). Assembly-based methods first reconstruct contigs from sequencing reads prior to downstream analysis, enabling genomic features to be interpreted within their broader genomic context. This can facilitate linkage of STs, serotypes, ARGs, and VGs to the same genomic fragment, which may be advantageous for strain-level characterisation and risk assessment (30, 31). However, assembly quality may be negatively affected by uneven coverage, repetitive genomic regions, and the coexistence of multiple closely related strains within a sample (32).

In contrast, read-based approaches analyse quality-controlled reads directly by mapping them to reference databases without an assembly step. Examples of databases used by the tools evaluated in this study include PubMLST, EcOH, CARD, and VFDB (33–36). Read-based methods are typically computationally efficient and avoid assembly-associated artefacts but may be limited by ambiguous read mapping and a reduced ability to reconstruct the broader genomic context (8, 37). Because both approaches present distinct advantages and limitations, inclusion of both workflows enabled a direct comparison of their effects on gene detection and strain typing.

For the read-based workflow, raw reads were processed using `fastp` (v1.0.1) for adapter trimming and quality filtering, followed by host read removal using `Bowtie2` (v2.5.2). Quality-controlled reads were subsequently supplied directly to the read-based analysis tools. For the assembly-based workflow, quality-controlled reads were assembled using MEGAHIT (v1.2.9) with parameters `-m 0.95, --min-contig-len 500, --min-count 2`, and `--k-list 21,41,61,81,99` (38). Preliminary analyses additionally evaluated metagenome binning using MetaBAT2 (v2.18) and DAS Tool (v1.1.3) (39, 40). However, binning consistently reduced recovery of the target genomic features and was therefore excluded from the final benchmarking analysis. Consequently, all assembly-based results reported in this study were generated from unbinned assemblies.

### 2.3 Tool Selection, Analysis and Evaluation

A panel of widely used bioinformatic tools was selected to represent major methodological approaches to genomic profiling. Tools were chosen based on their widespread use, open availability, applicability to *E. coli*, and representation of read-based, assembly-based, and hybrid analytical strategies. This diversity enabled a systematic comparison of how differing methodological assumptions influence performance when applied to standardized datasets. All tools were run using recommended default parameters. Tool versions and database releases are provided in Table 2.2.

**Table 2.2:**
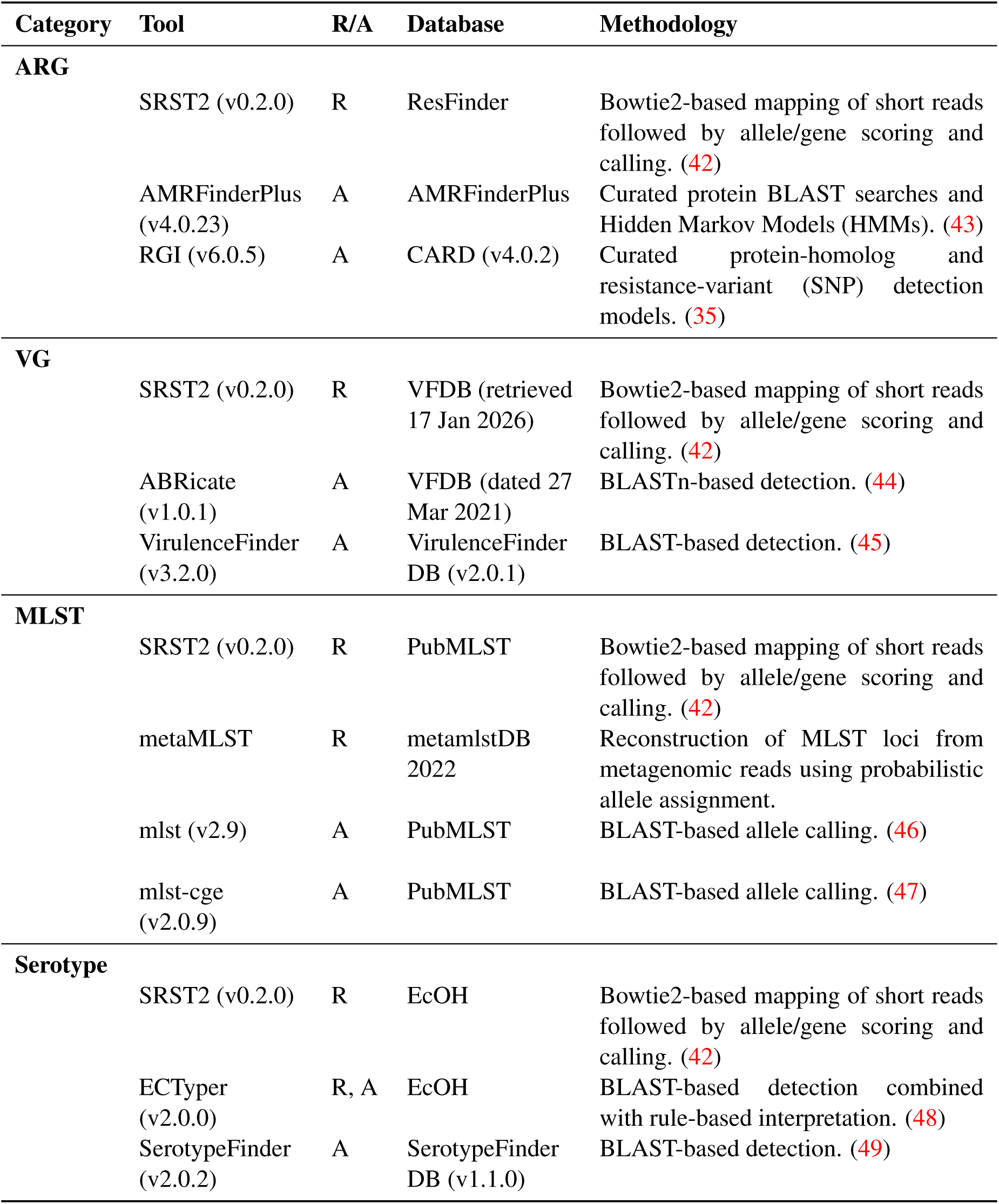
Comparison of the 10 bioinformatics tools used for genomic profiling of bacterial isolates, including antimicrobial resistance genes (ARGs), virulence genes (VG), multilocus sequence typing (MLST), and serotype prediction. For each tool, the analysis approach (R: read-based or A: assembly-based), reference database, and methodological framework are indicated. Bowtie2-based approaches rely on direct read mapping, whereas BLAST/BLASTn-based approaches perform sequence similarity searches against reference databases. HMMs identify conserved protein signatures, and probabilistic allele assignment infers the most likely MLST allelic profile from metagenomic reads. For SRST2, VFDB represents the DNA sequences of the full dataset, covering all genes related to known and predicted VFs in the VFDB database that was retrieved on 17 Jan 2026.

| Category | Tool | R/A | Database | Methodology |
| --- | --- | --- | --- | --- |
| **ARG** | SRST2 (v0.2.0) | R | ResFinder | Bowtie2-based mapping of short reads followed by allele/gene scoring and calling. (42) |
|  | AMRFinderPlus (v4.0.23) | A | AMRFinderPlus | Curated protein BLAST searches and Hidden Markov Models (HMMs). (43) |
|  | RGI (v6.0.5) | A | CARD (v4.0.2) | Curated protein-homolog and resistance-variant (SNP) detection models. (35) |
| **VG** | SRST2 (v0.2.0) | R | VFDB (retrieved 17 Jan 2026) | Bowtie2-based mapping of short reads followed by allele/gene scoring and calling. (42) |
|  | ABRicate (v1.0.1) | A | VFDB (dated 27 Mar 2021) | BLASTn-based detection. (44) |
|  | VirulenceFinder (v3.2.0) | A | VirulenceFinder DB (v2.0.1) | BLAST-based detection. (45) |
| **MLST** | SRST2 (v0.2.0) | R | PubMLST | Bowtie2-based mapping of short reads followed by allele/gene scoring and calling. (42) |
|  | metaMLST | R | metamlstDB 2022 | Reconstruction of MLST loci from metagenomic reads using probabilistic allele assignment. |
|  | mlst (v2.9) | A | PubMLST | BLAST-based allele calling. (46) |
|  | mlst-cge (v2.0.9) | A | PubMLST | BLAST-based allele calling. (47) |
| **Serotype** | SRST2 (v0.2.0) | R | EcOH | Bowtie2-based mapping of short reads followed by allele/gene scoring and calling. (42) |
|  | ECTyper (v2.0.0) | R, A | EcOH | BLAST-based detection combined with rule-based interpretation. (48) |
|  | SerotypeFinder (v2.0.2) | A | SerotypeFinder DB (v1.1.0) | BLAST-based detection. (49) |

ARG and VG presence was determined as a binary outcome (present/absent) for each gene of interest. MLST profiles were determined using the Achtman 7-gene scheme. Assembly-based methods generated a single ST prediction per assembly, while read-based methods could report up to two candidate STs per metagenome based on the paired-end read data. Serotypes were inferred based on O- and H-antigen gene detections.

Tool performance was evaluated by comparing predictions to PCR-derived ground truth. With simulated metagenomes containing multiple strains, the dominant strain was designated as the reference sequence type and serotype to be predicted. This choice reflects both the study objective and the behaviour of the evaluated tools, most of which are designed to report a single consensus strain type rather than reconstruct the complete strain composition of a metagenomic community. As a result, performance assessment focused on the ability of each workflow to recover the epidemiologically dominant strain profile from low-complexity metagenomes. Accuracy was calculated as the proportion of samples for which the predicted sequence type or serotype matched the PCR-validated label of the dominant strain. For gene detection, absence of a PCR annotation was not considered evidence of gene absence, as the reference annotations were derived from targeted PCR assays rather than comprehensive gene screening. Thus, gene performance assessment focused on recovery of PCR-confirmed targets rather than precision-based metrics requiring complete negative annotations. Gene recovery rates were calculated as recall, defined as the proportion of expected genes successfully detected by a tool (TP/(TP+FN)), where TP represents true positive detections and FN represents expected genes that were not recovered. This approach follows established benchmarking recommendations that advocate selecting performance metrics appropriate to the available ground truth and intended application (41). All metrics were calculated per simulated metagenome and averaged across samples.

## 3 Results

We first assessed the ability of read-based and assembly-based approaches to detect PCR-validated ARGs and VGs in Sections 3.1 and 3.2. We then evaluated recovery of strain-level epidemiological markers through MLST and serotype prediction in Sections 3.3 and 3.4. Because strain heterogeneity is expected to influence reconstruction of strain-specific features from metagenomic data, additional analyses examined the effect of increasing dominance of a single strain, ranging from heterogeneous multi-strain communities to single-strain metagenomes, on MLST and serotype prediction accuracys in Section 3.5. Finally, in Section 3.6, the potential benefit of long-read sequencing for strain typing in heterogeneous multi-strain metagenomes was assessed using simulated long-read datasets. Together, these analyses provide a comparative evaluation of gene detection and strain typing performance under different metagenomic scenarios.

### 3.1 ARG Recovery Exceeded 84% Across All Evaluated Tools

The ability of `SRST2, AMRFinderPlus`, and `RGI` to recover PCR-validated antimicrobial resistance genes was assessed using the ARG dataset comprising 10 *E. coli* metagenomes (34, 35, 43). Performance was evaluated using recovery of PCR-confirmed genes (Tables 3.1 and 3.2).

**Table 3.1:** Gene recovery rates per simulated metagenome for antibiotic resistance gene (ARG) and virulence gene (VG) detection across the assembly methods of AMRFinderPlus, RGI, ABRicate and VirulenceFinder. Gene recovery rate represents the proportion of expected genes successfully recovered in each sample and is reported as a percentage rounded to the nearest whole number. Mean gene recovery rate (MGRR) is reported per tool across the ten simulated samples and is shown to one decimal place. The poorest performing VG sample, VG03, is in bold.

| ARG |  |  | VG |  |  |
| --- | --- | --- | --- | --- | --- |
| Sample | AMRFinderPlus | RGI | Sample | ABRicate | VirulenceFinder |
| ARG01 | 5/7 (71%) | 5/7 (71%) | VG01 | 2/3 (67%) | 2/3 (67%) |
| ARG02 | 7/7 (100%) | 5/7 (71%) | VG02 | 2/3 (67%) | 2/3 (67%) |
| ARG03 | 7/7 (100%) | 7/7 (100%) | **VG03** | **1/3 (33%)** | **0/3 (0%)** |
| ARG04 | 6/7 (86%) | 6/7 (86%) | VG04 | 2/3 (67%) | 2/3 (67%) |
| ARG05 | 6/7 (86%) | 6/7 (86%) | VG05 | 2/3 (67%) | 2/3 (67%) |
| ARG06 | 6/7 (86%) | 6/7 (86%) | VG06 | 2/3 (67%) | 2/3 (67%) |
| ARG07 | 7/7 (100%) | 6/7 (86%) | VG07 | 2/3 (67%) | 2/3 (67%) |
| ARG08 | 7/7 (100%) | 6/7 (86%) | VG08 | 2/3 (67%) | 2/3 (67%) |
| ARG09 | 5/7 (71%) | 5/7 (71%) | VG09 | 2/3 (67%) | 2/3 (67%) |
| ARG10 | 7/7 (100%) | 7/7 (100%) | VG10 | 2/3 (67%) | 2/3 (67%) |
| MGRR | 90.0% | 84.3% | MGRR | 63.3% | 60.0% |

**Table 3.2:**
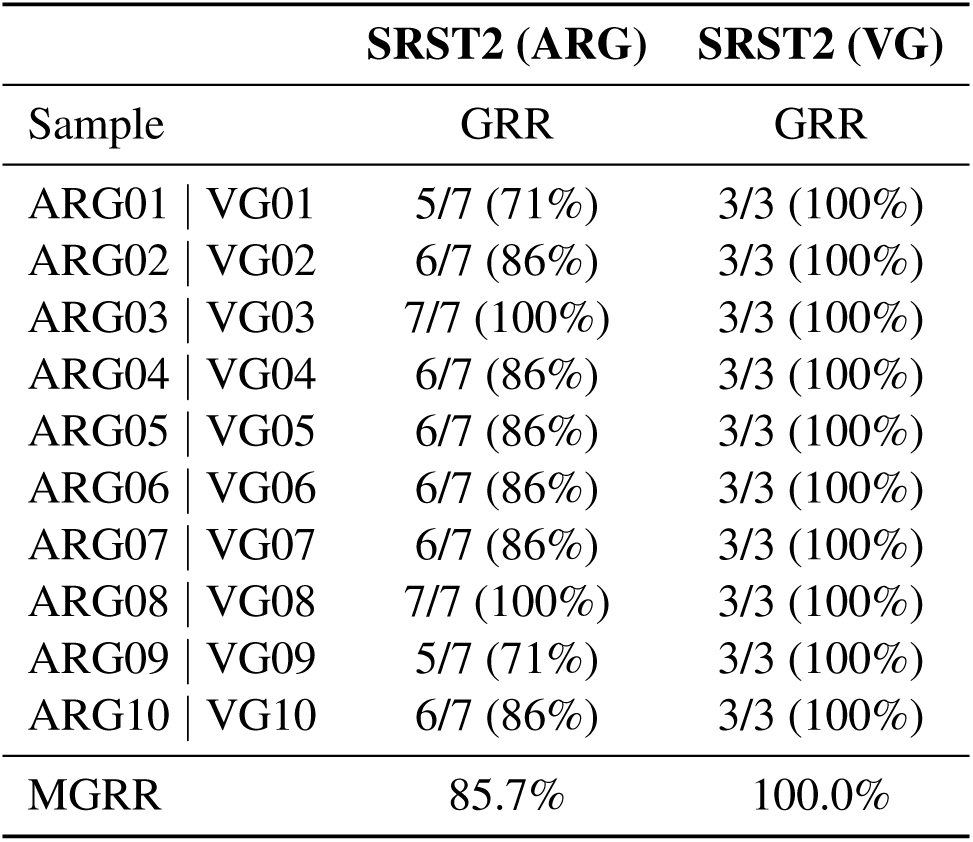
Gene recovery rates (GRRs) for antibiotic resistance gene (ARG) and virulence gene (VG) detection using read-based SRST2. Gene recovery rate represents the proportion of expected genes successfully recovered and is reported as a percentage rounded to the nearest whole number. Mean gene recovery rate (MGRR) averages the reported gene recovery rate per tool across all ten samples.

Overall, ARG recovery was high across all evaluated tools. `AMRFinderPlus` achieved the highest mean recovery rate (90.0%), followed by SRST2 (85.7%) and RGI (84.3%). Half of the simulated metagenomes saw complete recovery of all PCR-confirmed resistance determinants by one or more tools, indicating that clinically relevant ARGs can be reliably identified under low-complexity metagenomic conditions.

Differences between tools were generally small and largely attributable to the inconsistent recovery of a limited subset of genes. Across the evaluated samples, missed detections most frequently involved *sul3* and *tetD*, while the remaining genes of interest of *bla_TEM_*, *sul1*, *sul2*, *tetA*, and *tetB* were recovered consistently across most samples and tools. Database dependence was unlikely to explain these missed detections as all three methods successfully identified all seven ARGs in at least one sample, suggesting that their corresponding databases contain sufficient gene information to recover all the ARGs of interest. No resistance class was systematically missed, and reduced recovery rates were typically driven by failure to detect one or both of these less frequently recovered determinants rather than widespread loss of ARG detection capability.

### 3.2 Read-Based Analysis Achieved Complete Recovery of Virulence Genes

Virulence gene detection was evaluated using `SRST2, ABRicate`, and `VirulenceFinder` against PCR-derived annotations for *stx1*, *stx2* and *eae* (Tables 3.1 and 3.2) (34, 44, 45). SRST2 achieved the highest mean recovery rate at 100.0%, clearly outperforming the assembly-based approaches of `ABRicate` and VirulenceFinder that frequently recovered only a subset of the PCR-confirmed virulence genes.

The largest differences between methods were associated with incomplete recovery of *eae*. Verification of the databases used by these tools (VFDB used by `ABRicate`, VirulenceFinder DB) showed that *eae* was present in both (36, 45). The tool `ABRicate` recovered *eae* in only one sample, while VirulenceFinder failed to recover *eae* in all evaluated sample. In contrast, *stx2* was recovered consistently by both assembly-based tools (90.0%) and *stx1* showed intermediate recovery (80.0%). `ABRicate` reported subtypes *stx1a* and *stx1b* separately while VirulenceFinder reported family level *stx1* only. Since the PCR annotations stated *stx1*, both *stx1a* and *stx1b* were interpreted as positive hits to *stx1*. There was 100.0% concordance between `ABRicate` and VirulenceFinder recovery of *stx1* and *stx2* under these conditions. Both assembly tools experienced reduced performance for dataset VG03, which thus can be interpreted as the most challenging virulence sample in the benchmark.

### 3.3 No Tool Recovered Dominant Sequence Types from Multi-Strain Metagenomes

MLST prediction was evaluated using two assembly-based tools (mlst and mlst-cge) and two read-based tools (metaMLST and SRST2) (8, 34, 46, 47). Predictions were compared against the dominant PCR-validated sequence type present within each simulated metagenome (Table 3.3 and Supplementary Table S2).

**Table 3.3:** MLST predictions returned by assembly-based and read-based tools for simulated metagenomes. Predicted sequence types were compared against the dominant PCR-validated sequence type present in each metagenome, indicated in the column ‘Dominant ST’. All tools returned unassigned or incorrect sequence type calls for every sample. The output - from mlst indicated that the tool could not assign a valid ST from the MLST database. The output ‘NF’ from SRST2 indicated that the allele combination was not found. * next to the predicted ST indicated that there were mismatches against at least one of the alleles whileindicated that there was uncertainty in at least one of the alleles.

| Sample | `mlst` | `mlst-cge` | `metaMLST` | `SRST2` | Dominant ST |
| --- | --- | --- | --- | --- | --- |
| ST01 | - | Unknown | - | NF*? | 11 |
| ST02 | - | Unknown | - | NF*/NF*? | 17 |
| ST03 | - | Unknown | - | NF*?/NF? | 73 |
| ST04 | - | Unknown | - | NF*?/NF? | 12 |
| ST05 | - | Unknown | - | NF*? | 17 |
| ST06 | - | Unknown | - | NF*? | 43 |
| ST07 | - | Unknown | - | 7423*?/NF*? | 11 |
| ST08 | - | Unknown | - | NF*? | 73 |
| ST09 | - | Unknown | - | NF*? | 73 |
| ST10 | - | Unknown | - | NF*? | 73 |

The evaluated methods differ in their intended applications. metaMLST was specifically developed to reconstruct MLST profiles directly from metagenomic sequencing data and can report sequence types from mixed microbial communities (8). SRST2 was developed for read-based allele and sequence type assignment from short-read data and has been widely applied to both isolate genomes and metagenomic datasets (34). In contrast, mlst and mlst-cge are assembly-based approaches originally designed for typing individual bacterial isolates but are frequently applied to assemblies generated from metagenomic data (46, 47). Collectively, these tools therefore represent both methods explicitly developed for metagenomic strain typing and methods commonly used for sequence type inference from metagenome-derived assemblies.

In contrast to the high recovery rates observed for antibiotic resistance and virulence genes, none of the evaluated tools successfully recovered the dominant sequence type from the simulated multistrain metagenomes. Assembly-based tools (mlst and mlst-cge) and metaMLST returned only unassigned or unknown calls across all ten samples. SRST2 produced one ambiguous sequence type prediction of 7423 that did not match the expected dominant sequence type of 11, resulting in an overall accuracy of 0% for all tools. Dominant sequence types represented within the dataset included ST11, ST12, ST17, ST43, ST73, all of which are previously described sequence types within the Achtman MLST scheme (28). Despite this, no tool recovered a valid sequence type assignment. Notably, this lack of recovery was observed even for metaMLST, which was specifically designed for MLST profiling of metagenomic datasets.

### 3.4 SerotypeFinder Provided the Most Accurate Serotype Reconstruction

Four serotype prediction workflows were evaluated using the simulated metagenomic dataset: assembly-based ECTyper and SerotypeFinder, and read-based ECTyper and SRST2. Compared with antimicrobial resistance and virulence gene detection, serotype prediction showed substantially lower performance across all methods. Yet, when compared against MLST, serotype prediction proved much more successful.

Performance was assessed at both the antigen and serotype levels (Table 3.4). O-antigen recovery and H-antigen recovery were calculated independently as the proportion of the expected antigens recovered. Overall antigen recovery was calculated as the proportion of all expected antigens (O and H combined) recovered across samples. Complete serotype recovery required successful identification of both the expected O and H antigens within the same sample. This approach enabled differentiation between partial serotype assignments and complete serotype reconstruction.

**Table 3.4:** Performance of serotype prediction tools. O-antigen recovery and H-antigen recovery were calculated as the proportion of expected antigens recovered. Overall antigen recovery was calculated as the proportion of all expected antigens (O and H combined) recovered across samples. Complete serotype recovery required recovery of both the expected O and H antigens within a sample. The mean recovery for each of these four performance metrics across the four different tools was summarised.

| Tool | O-antigen | H-antigen | Overall antigen recovery | Complete serotype recovery |
| --- | --- | --- | --- | --- |
| Assembly-based |  |  |  |  |
| **ECTyper** | 8/10 (80.0%) | 6/10 (60.0%) | 14/20 (70.0%) | 5/10 (50.0%) |
| **SerotypeFinder** | 8/10 (80.0%) | 9/10 (90.0%) | 17/20 (85.0%) | 8/10 (80.0%) |
| Read-based |  |  |  |  |
| **ECTyper** | 4/10 (40.0%) | 0/10 (0.0%) | 4/20 (20.0%) | 0/10 (0.0%) |
| **SRST2** | 6/10 (60.0%) | 6/10 (60.0%) | 12/20 (60.0%) | 3/10 (30.0%) |
| Mean recovery | 65.0% | 52.5% | 58.8% | 40.0% |

Performance varied considerably among methods. The assembly-based approaches reported at least 70% overall antigen recovery, with SerotypeFinder outperforming ECTyper. Read-based ECTyper was the least effective at 20.0% overall antigen recovery while SRST2 showed intermediate antigen recovery at 60.0%.

Recovery of individual antigens was generally more common than complete serotype reconstruction. Although one antigen was often correctly identified, failure to recover both the O- and H-antigen within the same sample substantially reduced complete serotype recovery across all methods. Differences between O-antigen and H-antigen recovery were also evident. O-antigen recovery was relatively robust across most workflows (40.0–80.0%), while H-antigen recovery showed substantially greater variability between tools (0.0–90.0%). Notably, SerotypeFinder demonstrated particularly strong recovery of both antigens, resulting in the highest rate of complete serotype reconstruction among the evaluated approaches at 80%.

Complete serotype recovery was achieved in at least one sample by three of the four evaluated methods. The highest level of complete serotype recovery was reported by SerotypeFinder, while read-based ECTyper did not recover a complete serotype in any of the evaluated samples.

### 3.5 Reduced Strain Complexity Improved MLST and Serotype Prediction

The complete failure of MLST recovery and the reduced success of serotype prediction in the multi-strain datasets suggested that strain heterogeneity may be a major factor limiting strain-level characterisation. To investigate the impact of strain heterogeneity on strain typing performance, a series of metagenomic datasets was simulated in which a dominant sequence type was present at fixed relative abundances. A single-strain dataset was represented by having a single sequence type at 100% abundance, while the four dominant-strain datasets contained a dominant sequence type at 60%, 70%, 80% and 90% relative abundance respectively. These datasets were compared with the original multi-strain simulations that had been generated using log-normal abundance distributions, with results reported in Section 3.3. The distribution of dominant STs and their abundances across all six datasets can be found in Supplementary Figure S1. The dominant ST abundance in the multi-strain dataset ranged from 33% to 88%, with many simulations comprising multiple sequence types contributing a substantial fraction of the total sample. This design enabled evaluation of tool performance across a gradient of decreasing strain complexity.

MLST prediction accuracy improved consistently with increasing dominance of a single strain and thus decreasing complexity in the sample (Figure 3.1). Assembly-based methods exhibited the greatest improvement as strain heterogeneity reduced. Both mlst and mlst-cge achieved only limited accuracy when the dominant sequence type represented 60–80% of the metagenome, but performance increased substantially at higher abundances, reaching 90.0% accuracy in the single-strain datasets. Among the assembly-based methods, mlst-cge showed slightly higher performance when faced with more heterogenous metagenomes, though both methods converged on identical accuracy at 90% and 100% abundance.

**Figure 3.1:**
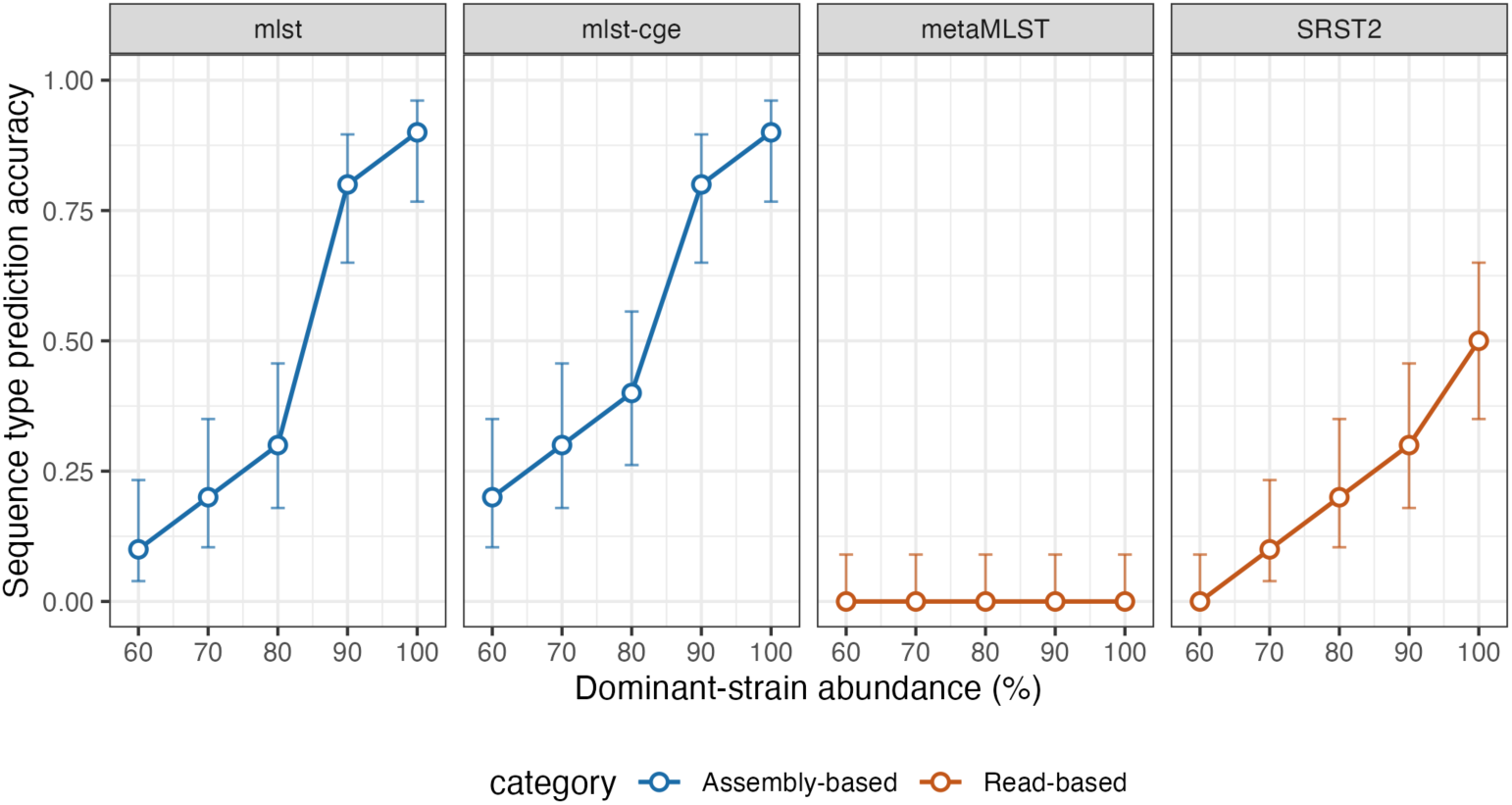
Sequence type (ST) prediction accuracy of four multi-locus sequence typing (MLST) calling tools across a gradient of dominant-strain relative abundance in simulated metagenomes. Assembly-based tools (mlst, mlst-cge) and read-based tools (metaMLST, SRST2) are shown in separate panels, coloured by category. For each abundance level (60–100%), accuracy is calculated as the proportion of correct ST calls across 10 independently simulated metagenomes. A call was scored as correct if it matched the true dominant ST, or, for read-based methods, if the true dominant ST was reported from at least one of the paired-end reads. Error bars show the 68% Wilson score confidence interval (about one standard deviation) for the estimated proportion at each point.

SRST2 also demonstrated improved performance as dominant-strain abundance increased, although accuracy remained below that of the assembly-based approaches across all abundance levels. Accuracy increased from 0.0% at 60% abundance to 50.0% in single-strain metagenomes. In contrast, metaMLST failed to recover any correct sequence type assignments across the entire abundance gradient.

The original multi-strain datasets generated using log-normal abundance distributions represented the most challenging scenario. Under these conditions, no tool successfully recovered the correct sequence type. Although confidence intervals overlapped between some neighbouring abundance levels, the overall trend of improved sequence type recovery was consistent across all successful methods as the datasets approached single-strain composition.

Similarly, in serotyping, a single-strain dataset was created in which a single serotype represented 100% of the metagenome, while dominant-strain datasets contained a serotype at 60%, 70%, 80% and 90% relative abundance. They were compared with the original multi-strain dataset that had been generated using log-normal abundance distributions, with results reported in Section 3.4. The distribution of dominant serotypes and their abundances across all six datasets can be found in Supplementary Figure S2.

Serotype prediction performance was likewise strongly influenced by strain composition (Figure 3.2, Supplementary Tables S3-S6). Across all methods, recovery rates generally improved as the abundance of the dominant serotype increased, although the relationship was not strictly monotonic across intermediate (70–90%) abundance levels.

**Figure 3.2:**
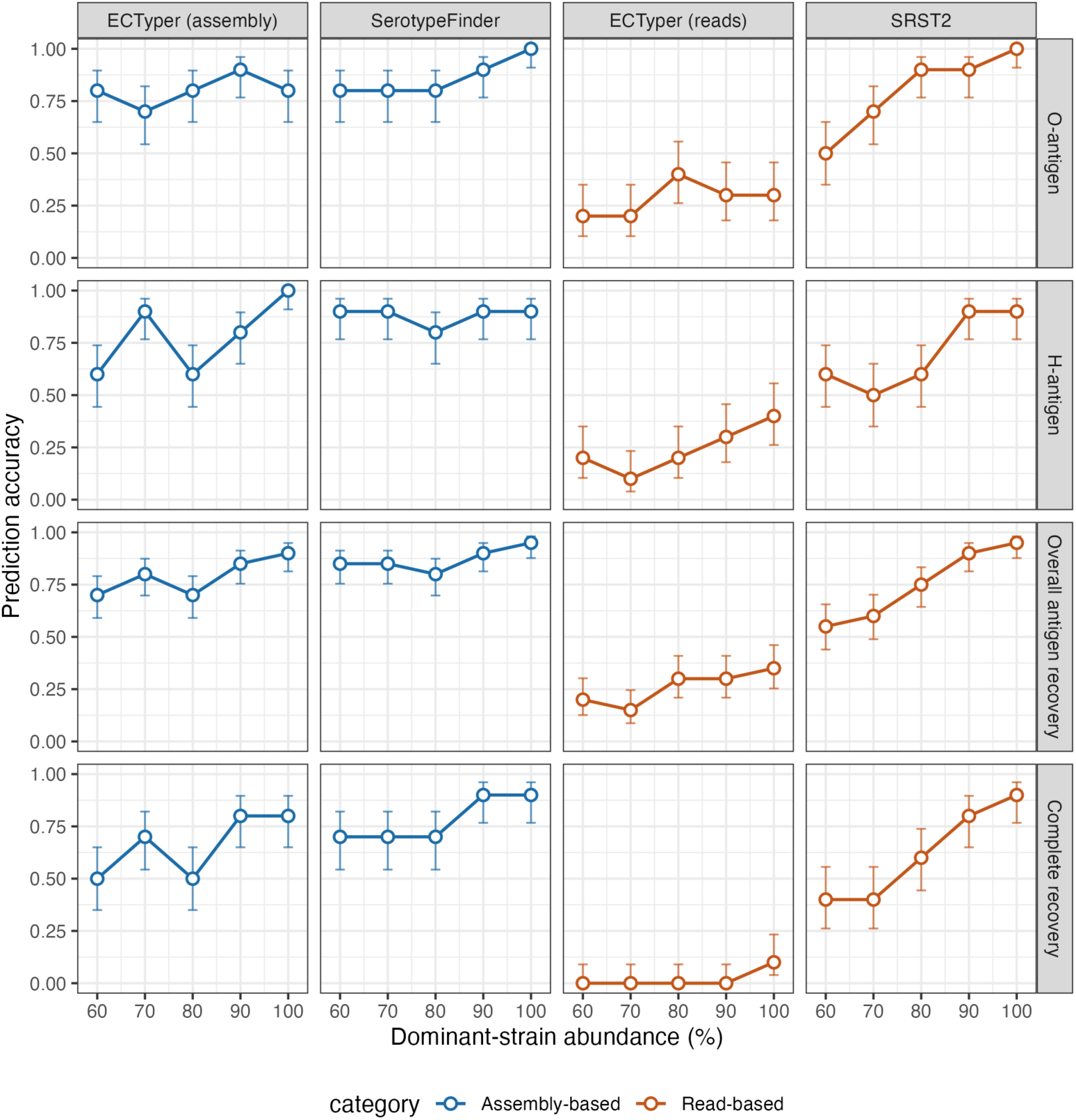
Serotype (O- and H-antigen) prediction accuracy of four serotyping tools across a gradient of dominant-strain relative abundance in simulated metagenomes. Columns show assembly-based tools (ECTyper, SerotypeFinder) and read-based tools (ECTyper, SRST2); rows show, from top to bottom, O-antigen recovery, H-antigen recovery, overall antigen recovery (O- and H-antigen calls pooled), and complete serotype recovery (both O and H antigens correctly identified in the same simulation). For each abundance level (60–100%), accuracy is the proportion of correct calls across 10 independently simulated metagenomes, where a call was scored correct if the true antigen appeared among the antigens reported for an ambiguous multi-allele call. Error bars show the 68% Wilson score confidence interval (about one standard deviation) for the estimated proportion (n = 10 per point for O-antigen, H-antigen and complete recovery; n = 20 pooled O- and H-antigen calls per point for overall antigen recovery).

Assembly-based methods consistently achieved the highest recovery rates. SerotypeFinder exhibited the strongest overall performance across the abundance gradient, with overall antigen recovery ranging from 85.0% in multi-strain log-normal metagenomes to 95.0% in single-strain metagenomes. Assembly-based ECTyper showed a similar pattern but achieved lower recovery rates in the more heterogeneous datasets, with overall antigen recovery declining from 90.0% in single-strain metagenomes to 70.0% in the multi-strain log-normal samples. Read-based SRST2 performed comparably to the assembly-based tools under low-complexity conditions but exhibited a larger reduction in recovery as strain heterogeneity increased. Read-based ECTyper consistently showed the lowest overall performance across all dataset compositions. Because both assembly-based serotyping tools operated on identical assemblies yet produced different recovery rates, the observed better performance of SerotypeFinder and assembly-based ECTyper cannot be attributed solely to assembly quality.

The effect of strain composition differed markedly between O-antigen and H-antigen recovery. O-antigen recovery remained relatively stable across abundance levels, particularly for the assembly-based tools, with recovery rates typically varying by 10-20 percentage points across the entire gradient. In contrast, H-antigen recovery showed substantially greater sensitivity to strain composition. For assembly-based ECTyper, H-antigen recovery declined from 100.0% in single-strain metagenomes to 60.0% in multi-strain communities, while read-based ECTyper recovered no H-antigens in the multi-strain dataset and only 40.0% in the single-strain samples.

The superior performance of SerotypeFinder was attributable largely to improved H-antigen recovery. In the multi-strain dataset, both assembly-based tools recovered O-antigens at identical rates (80.0%), yet SerotypeFinder achieved substantially higher H-antigen recovery (90.0% versus 60.0%) and consequently higher complete serotype recovery (80.0% versus 50.0%) than assembly-based ECTyper. A similar pattern was observed across several dominant-abundance datasets, suggesting that robust H-antigen detection represents a key determinant of successful serotype reconstruction.

Differences among tools became more pronounced when complete serotype recovery was considered. Although several tools achieved moderate recovery of individual antigens, successful reconstruction of both O- and H-antigens in the same sample remained substantially more challenging. The highest complete serotype recovery was consistently achieved by SerotypeFinder, increasing from 80.0% in multi-strain communities to 90.0% in single-strain metagenomes. SRST2 and assembly-based ECTyper showed comparable trends but achieved lower recovery as heterogeneity increased.

Overall, these results identify strain heterogeneity as a major determinant of serotype prediction performance in metagenomic datasets. Performance generally improved as communities became increasingly dominated by a single serotype, but successful serotype reconstruction remained dependent on recovery of both antigen-associated loci.

### 3.6 Long-Read Sequencing Did Not Improve Strain Typing Recovery

To evaluate if longer sequencing reads could improve MLST and serotype prediction, long reads were simulated using Badread (v0.4.1) (50). Long-read simulation was performed for the log-normal multi-strain abundance dataset only, at a target length of 15,000 bp using Badread’s pacbio2016 error and quality-score model. This was intended to represent the most challenging and realistic composition for evaluating whether increased read length could compensate for the difficulty short reads showed in this dataset category.

Sequence type was predicted directly from uncorrected long reads using Krocus (v1.0.3), which infers the closest-matching allele for each of the seven Achtman MLST genes from k-mer coverage without requiring assembly (51). Serotype prediction was performed using ECTyper (v2.0.0) in its native raw-read mode, which aligns reads against curated O- and H-antigen gene databases to build a consensus sequence prior to typing (48).

Both tools recovered no sequence types and no serotypes, compared to the more positive performance of the serotyping tools on the short-read multi-strain dataset.

## 4 Discussion

In this study, we established a controlled benchmarking framework to evaluate the performance of widely used genomic profiling tools under low-complexity metagenomic conditions representative of enrichment-based surveillance workflows (52, 53). By leveraging PCR-validated annotations as an independent ground truth, we directly assessed the accuracy of tools across multiple clinically relevant feature classes, including antibiotic resistance genes (ARGs), virulence genes (VGs), MLST and serotype.

### 4.1 Gene Detection

#### Gene detection is more resilient to assembly fragmentation than strain-level markers

A key finding was that tool performance varied substantially according to the genomic feature being investigated. ARG detection was the most robust analytical task, with all evaluated tools achieving high gene recovery rates exceeding 84.3% (Tables 3.1 and 3.2) (35, 54). The relatively small differences between AMRFinderPlus, RGI, and SRST2 suggest that recovery of the targeted resistance determinants was not strongly limited by assembly quality. Resistance genes are often identified through sequence similarity to curated reference databases rather than through reconstruction of broader genomic context (35, 43). The differences observed between methods indicate that resistance loci are generally well represented within both assembled contigs and raw read datasets. As a result, successful recovery of individual ARGs may depend less strongly on the assembler performance than analyses requiring linkage of multiple loci within the same genome. These results are encouraging for surveillance applications where rapid identification of clinically relevant resistance determinants is often the primary objective (35, 43).

The reduced recovery observed for a small subset of ARGs likely reflects characteristics of individual loci rather than systematic limitations of any specific tool. Missed ARG detections were primarily associated with *sul3* and *tetD*. Several factors may explain this pattern. First, both genes are less commonly reported in *E. coli* than widely distributed determinants such as *sul1*, *sul2*, *tetA*, and *tetB*, potentially resulting in fewer representative reference sequences and reduced detection robustness across bioinformatic databases and tools (35). Although *sul3* is generally reported at lower prevalence than *sul1* and *sul2*, the gene was successfully detected in at least one sample by all evaluated tools, indicating that it was represented within the underlying reference databases (55, 56). Likewise, because *sul1*, *sul2*, and *sul3* are all commonly associated with mobile genetic elements, differences in recovery cannot be attributed solely to plasmid association. Instead, the reduced recovery may reflect a combination of factors including sequence diversity among *sul3* alleles, variation in local genomic context, incomplete recovery of resistance-bearing regions, or interactions between these factors and tool-specific detection thresholds. Second, both assembly-based and read-based approaches rely on sufficient sequence coverage and similarity to reference sequences. Sequence divergence, fragmented assembly of resistance-bearing regions, or partial gene recovery may therefore reduce confidence in gene assignment even when the corresponding resistance determinant is present in the metagenome. Because missed detections were largely restricted to *sul3* and *tetD*, while recovery of *bla_TEM_, sul1, sul2, tetA*, and *tetB* remained consistently high, the reduced performance appears to more likely reflect characteristics of these specific genes and their genomic contexts than systematic shortcomings of any individual tool.

#### Virulence gene recovery was influenced by gene-specific biological characteristics

In contrast to ARG detection, VG recovery showed a stronger dependence on analytical approach. Read-based SRST2 recovered the targeted VGs more consistently than the assembly-based methods (Tables 3.1 and 3.2). This pattern suggests that successful recovery of virulence determinants may depend more strongly on reconstruction of the genomic regions in which these genes occur.

Reduced VG recovery was driven primarily by failure to detect *eae*, while *stx1* and *stx2* were recovered consistently across methods. The *eae* gene encodes intimin, a surface-exposed adhesin involved in host attachment, and comprises numerous variants that exhibit substantial nucleotide sequence divergence (57). Involved in direct host-pathogen interactions, *eae* is likely subject to stronger diversifying selection than the other genes examined in this study. Divergence between recovered *eae* sequences and reference database entries may reduce annotation sensitivity, particularly for assembly-based approaches that rely on high sequence similarity for gene assignment.

An additional explanation for the observed difference between ARG and VG recovery may lie in the interaction between sequence diversity and assembly-based analysis. Assembly-based workflows depend on successful reconstruction of a representative consensus sequence before database comparison, while read-based approaches can identify matching sequence signatures directly from individual reads. As a result, sequence divergence within *eae* may have had a greater impact on assembly-based VG recovery than on the more conserved ARG targets evaluated in this study.

The *eae* gene occurs within the locus of enterocyte effacement (LEE), a horizontally acquired pathogenicity island containing numerous virulence-associated genes and regulatory elements (58–60). Although the genomic architecture of the LEE island may also influence recovery in assembly-based workflows, successful detection ultimately depends on recognition of the *eae* sequence itself.

Taken together, these observations suggest that recovery of virulence genes is influenced not only by the analytical method used but also by the biological characteristics of the target gene. Recovery of ARGs appeared comparatively robust across analytical strategies, while VG detection was more strongly influenced by locus-specific biological characteristics, including sequence diversity and potentially the genomic context in which virulence genes occur. The reduced recovery of *eae* relative to *stx1* and *stx2* suggests that variation within target genes themselves may be at least as important as assembly-related factors in determining detection success.

### 4.2 Strain Typing

#### Strain heterogeneity was a dominant factor limiting strain typing

The most notable finding of this study was the marked contrast between gene detection and strain-level characterisation. While ARGs and VGs were recovered with moderate to high success across most samples, recovery of sequence types and serotypes was substantially more limited. This difference likely reflects the fundamentally different information requirements of these analyses. Identification of individual genes requires only local evidence for gene presence, whereas MLST and serotyping require reconstruction of multiple loci originating from the same strain. In mixed-strain metagenomes, alleles and antigen-associated loci derived from different genomes may coexist within the same sample, increasing ambiguity during assembly and downstream typing (42, 47, 61).

The strain-composition experiments suggest that strain heterogeneity was a major factor influencing typing performance. As the abundance of the dominant strain increased, both MLST and serotype prediction accuracy improved across several workflows (Figures 3.1 and 3.2). Substantial improvements in MLST and serotype recovery became apparent once the dominant strain exceeded approximately 80–90% relative abundance, suggesting that successful strain typing may require near-clonal enrichment of target populations. This pattern is consistent with reduced competition among strain-specific alleles and antigen loci, allowing more coherent reconstruction of strain-level profiles. Similar challenges have been reported in strain-resolved metagenomic studies, where the presence of multiple closely related strains complicates assignment of genomic features to individual genomes (62). With metagenomes being fundamentally heterogenous, the observed decline in typing performance highlights a key limitation of current approaches in reconstructing a complete and internally consistent strain profile when applied to realistic surveillance datasets containing multiple closely related strains.

The MLST results are consistent with this interpretation. MLST schemes require recovery of a complete allelic profile across multiple housekeeping genes, and ambiguity at a single locus can prevent sequence type assignment even when most loci are successfully recovered (13, 28). Thus, partial recovery of MLST loci may provide insufficient information for confident assignment in mixed-strain communities. The improvement observed in low-complexity datasets suggests that successful MLST prediction depends strongly on the ability to associate alleles with a single dominant genome. Although typing accuracy improved markedly as strain heterogeneity decreased, several methods failed to achieve complete recovery even in the single-strain dataset. This observation suggests that strain heterogeneity was not the sole factor limiting performance. The poor performance of metaMLST, a tool specifically developed for MLST reconstruction from metagenomic sequence data, further supports this theory that the challenge extends beyond simple strain heterogeneity. The discrepancy may reflect differences between the assumptions underlying the algorithm and the characteristics of the simulated datasets, highlighting the importance of validating specialised metagenomic typing tools under diverse conditions rather than assuming superior performance based on intended application. Additionally, residual errors likely arose from methodological limitations inherent to the typing workflows themselves, including incomplete recovery of MLST loci, ambiguous allele assignments, assembly fragmentation or database-dependent classification thresholds. Consequently, removal of strain heterogeneity substantially improved typing accuracy but was not sufficient to guarantee correct strain type assignment in every sample in the single-strain dataset.

A similar pattern was observed for serotype prediction. The recovery of individual antigen-associated loci was generally more successful than recovery of complete serotypes, indicating that reconstruction of a coherent antigen profile represents a greater challenge than detection of individual loci (Table 3.4). This finding mirrors the MLST results and suggests in mixed-strain metagenomes, successful linkage of multiple strain-specific features represents a major challenge beyond simple locus detection. Previous studies have similarly reported that metagenomic strain typing becomes increasingly difficult when closely related strains contribute overlapping genomic signals (42, 47, 61). The persistence of serotyping errors in single-strain metagenomes further supports this interpretation. Correct serotype assignment requires simultaneous recovery of both O- and H-antigen determinants, and incomplete recovery of either locus can prevent accurate classification. Thus, even under conditions where strain heterogeneity was effectively eliminated, technical limitations associated with locus reconstruction and classification remained evident.

#### Long reads did not overcome the challenge, suggesting that the bottleneck is strain deconvolution not sequence continuity

An additional observation was that increased read length alone did not improve strain typing performance. Neither Krocus nor read-based ECTyper recovered sequence types or serotypes from the simulated long-read datasets despite operating directly on raw reads. While long reads can improve genome contiguity under some circumstances, the present results suggest that sequence continuity was not the primary limitation in these datasets. Instead, the coexistence of multiple closely related strains may have generated sufficient allelic and antigenic ambiguity to prevent confident assignment even when substantially longer reads were available. This interpretation is consistent with the strain-composition experiments, which indicate that strain complexity had a greater influence on typing success than sequencing strategy itself.

Taken together, these findings suggest that the principal obstacle to metagenomic strain typing is reconstruction of coherent strain-level profiles in the presence of multiple closely related genomes. In contrast to gene detection, which relies primarily on identifying individual loci, MLST and serotyping require accurate association of multiple genomic features with the same biological strain. Current isolate-oriented typing methods thus appear best suited to metagenomes dominated by a single strain, while more heterogeneous communities will likely require dedicated strain-resolved approaches.

### 4.3 Implications and Limitations

The present results have important implications for surveillance applications. Although serotype-associated antigens could often be detected individually, complete serotype assignments and sequence type predictions were considerably less reliable than ARG or virulence gene detection, particularly in datasets containing multiple strains. Therefore, strain-level epidemiological characterisation derived from metagenomic data should be interpreted cautiously and regarded as supportive rather than definitive evidence unless corroborated by isolate sequencing, strain-resolved assembly methods, or complementary epidemiological information.

Several limitations should be considered. First, all simulations consisted exclusively of *E. coli*and therefore do not capture the taxonomic complexity of environmental or wastewater metagenomes (6). Second, metagenome binning was evaluated but reduced recovery of ARGs, VGs, MLST profiles, and serotypes relative to direct analysis of assembled contigs, likely reflecting the difficulty of binning closely related strains. As a result, only unbinned assemblies were retained for benchmarking. Third, the study evaluated a limited set of clinically relevant genes and typing targets, and additional datasets will be needed to assess broader generalisability. Lastly, because ground-truth information was available only for validated targets, gene detection was assessed using recovery of expected genes rather than precision-based measures.

Despite these limitations, the benchmark provides a controlled assessment of commonly used genomic profiling approaches in mixed-strain *E. coli* metagenomes. The exclusive use of single-species metagenomes should be interpreted as a reductionist framework intended to isolate the effects of strain heterogeneity. In environmental surveillance samples, strain heterogeneity occurs simultaneously with taxonomic complexity, variable abundance distributions, and background DNA from numerous species. Consequently, the levels of strain-typing performance reported here may represent an upper bound on what can be expected in real wastewater metagenomes. Furthermore, the results suggest that detection of epidemiologically relevant genes is achievable under these conditions, while strain-level characterisation remains substantially more challenging. Future developments in strain-resolved metagenomics, hybrid and long-read sequencing, and haplotype-aware analytical methods may improve recovery of MLST and serotype information from complex microbial communities (62).

## 5 Conclusion

This study established a controlled benchmarking framework to evaluate assembly-based and read-based bioinformatic tools for genomic profiling of *E. coli* in low-complexity metagenomic datasets using PCR-validated ground truth. By assessing ARGs, VGs, STs and serotypes across multiple analytical approaches, the study examined how analytical performance varies across different levels of genomic characterisation.

Tool performance was highly dependent on the genomic feature being investigated. ARG detection was consistently robust across tools, while VG recovery showed greater sensitivity to analytical approach. In contrast, strain-level characterisation proved substantially more challenging. Recovery of individual loci was often possible, but reconstruction of complete sequence type and serotype profiles was substantially reduced in more complex mixed-strain communities. The abundance-gradient experiments further indicated that strain heterogeneity was a major factor influencing typing performance, while the long-read analyses suggested that increased read length was insufficient to overcome the challenges associated with complex strain mixtures.

These findings highlight an important distinction between gene detection and strain typing in metagenomic analyses. Detection of individual resistance and virulence determinants can be achieved reliably under low-complexity conditions, while reconstruction of coherent strain-level profiles remains substantially more demanding when multiple closely related strains are present. Thus, the reliability of metagenomic surveillance depends significantly on the biological question being addressed. However, identification of individual genes represents only one level of genomic surveillance. In many epidemiological and public health applications, the critical question is not simply whether resistance or virulence determinants are present, but whether they co-occur within the same organism and can be linked to a particular strain background. Determining which ARGs and virulence genes are carried by the same genome, and subsequently associating those features with specific sequence types or serotypes, requires a level of genomic resolution similar to that needed for strain typing itself. The present findings therefore suggest that while gene detection is achievable under low-complexity metagenomic conditions, reconstruction of biologically meaningful genotype combinations remains a substantially greater challenge.

As metagenomic surveillance becomes increasingly integrated into public health, veterinary, and environmental monitoring programmes, careful interpretation of strain-level classifications will remain important. While current approaches are suitable for surveillance objectives focused on antimicrobial resistance and virulence determinants, strain-level classifications such as MLST and serotype should be interpreted cautiously unless supported by complementary evidence. Through future developments in strain-resolved metagenomics, recovery of epidemiologically relevant strain characteristics from mixed microbial communities may be improved. More importantly, such approaches may enable reliable linkage of resistance and virulence genes, sequence types, and serotypes within individual genomes, providing the level of resolution required for next-generation genomic surveillance.

## Data and Code Availability

All simulated metagenomes have been deposited in Figshare at https://doi.org/10.6084/m9.figshare.33940657. All generated results and code supporting the findings of this study have been deposited in GitHub at https://github.com/dibsondivya/MixEcoliBench under the MIT License. The associated README contains more detailed information.

## Acknowledgments

We acknowledge the National Academic Infrastructure for Supercomputing in Sweden (NAISS), partially funded by the Swedish Research Council through grant agreement no. 2022-06725, for awarding this project access to NSC Tetralith for the performing of computations. This work was supported by the Swedish Research Council through grant agreement no. 2024-03949, and eSSENCE, a Swedish strategic research programme in e-Science.

## References

1. Grassly NC, Shaw AG, Owusu M. 2025. Global wastewater surveillance for pathogens with pandemic potential: opportunities and challenges. Lancet Microbe 6:100939. 10.1016/j.lanmic.2024.07.002.

2. Parkins MD, Lee BE, Acosta N, Bautista M, Hubert CRJ, Hrudey SE, Frankowski K, Pang XL. 2024. Wastewater-based surveillance as a tool for public health action: SARS-CoV-2 and beyond. Clin Microbiol Rev 37:e00103–22. 10.1128/cmr.00103-22.

3. Philo SE, De León KB, Noble RT, Zhou NA, Alghafri R, Bar-Or I, Darling A, D’Souza N, Hachimi O, Kaya D, Kim S, Gaardbo Kuhn K, Layton BA, Mansfeldt C, Oceguera B, Radniecki TS, Ram JL, Saunders LP, Shrestha A, Stadler LB, Steele JA, Stevenson BS, Vogel JR, Bibby K, Boehm AB, Halden RU, Delgado Vela J. 2024. Wastewater surveillance for bacterial targets: current challenges and future goals. Appl Environ Microbiol 90:e01428–23. 10.1128/aem.01428-23.

4. MacGowan A, Macnaughton E. 2017. Antibiotic resistance. Medicine (Abingdon) 45:622–628. 10.1016/j.mpmed.2017.07.006.

5. Antimicrobial Resistance Collaborators. 2022. Global burden of bacterial antimicrobial resistance in 2019: a systematic analysis. Lancet 399:629–655. 10.1016/S0140-6736(21)02724-0.

6. Hendriksen RS, Munk P, Njage P, van Bunnik B, McNally L, Lukjancenko O, Röder T, Nieuwenhuijse D, Pedersen SK, Kjeldgaard J, Kaas RS, Clausen PTLC, Vogt JK, Leekitcharoenphon P, van de Schans MGM, Zuidema T, de Roda Husman AM, Rasmussen S, Petersen B, Global Sewage Surveillance project consortium, Amid C, Cochrane G, Sicheritz-Ponten T, Schmitt H, Alvarez JRM, Aidara-Kane A, Pamp SJ, Lund O, Hald T, Woolhouse M, Koopmans MP, Vigre H, Petersen TN, Aarestrup FM. 2019. Global monitoring of antimicrobial resistance based on metagenomics analyses of urban sewage. Nat Commun 10:1124. 10.1038/s41467-019-08853-3.

7. Noman SM, Shafiq M, Bibi S, Mittal B, Yuan Y, Zeng M, Li X, Olawale OA, Jiao X, Irshad M. 2023. Exploring antibiotic resistance genes, mobile gene elements, and virulence gene factors in an urban freshwater samples using metagenomic analysis. Environ Sci Pollut Res Int 30:2977–2990. 10.1007/s11356-022-22197-4.

8. Zolfo M, Tett A, Jousson O, Donati C, Segata N. 2017. MetaMLST: multi-locus strain-level bacterial typing from metagenomic samples. Nucleic Acids Res 45:e7. 10.1093/nar/gkw837.

9. Chen Z, Grim CJ, Ramachandran P, Meng J. 2024. Advancing metagenome-assembled genome-based pathogen identification: unraveling the power of long-read assembly algorithms in Oxford Nanopore sequencing. Microbiol Spectr 12:e00117–24. 10.1128/spectrum.00117-24.

10. Gholipour S, Lee D, Tokuriki N. 2025. Molecular Evolution and Origins of Antibiotic Resistance Genes. Biochemistry 64:2516–2528. 10.1021/acs.biochem.5c00044.

11. Wassenaar TM, Gaastra W. 2001. Bacterial virulence: can we draw the line? FEMS Microbiol Lett 201:1–7. 10.1111/j.1574-6968.2001.tb10724.x.

12. López-Banda DA, Carrillo-Casas EM, Leyva-Leyva M, Orozco-Hoyuela G, Manjarrez-Hernández Á H, Arroyo-Escalante S, Moncada-Barrón D, Villanueva-Recillas S, Xicohtencatl-Cortes J, Hernández-Castro R. 2014. Identification of virulence factors genes in *Escherichia coli* isolates from women with urinary tract infection in Mexico. BioMed Res Int 2014:959206. 10.1155/2014/959206.

13. Maiden MC, Bygraves JA, Feil E, Morelli G, Russell JE, Urwin R, Zhang Q, Zhou J, Zurth K, Caugant DA, Feavers IM, Achtman M, Spratt BG. 1998. Multilocus sequence typing: a portable approach to the identification of clones within populations of pathogenic microorganisms. Proc Natl Acad Sci U S A 95:3140–3145. 10.1073/pnas.95.6.3140.

14. Ørskov F, Ørskov I. 1984. Serotyping of *Escherichia coli*, p 43–112. *In* Bergan T (ed), Methods in microbiology, vol 14. Academic Press, London, United Kingdom. 10.1016/S0580-9517(08)70447-1.

15. Alhadlaq MA, Aljurayyad OI, Almansour A, Al-Akeel SI, Alzahrani KO, Alsalman SA, Yahya R, Al-Hindi RR, Hakami MA, Alshahrani SD, Alhumeed NA, Al Moneea AM, Al-Seghayer MS, AlHarbi AL, Al-Reshoodi FM, Alajel S. 2024. Overview of pathogenic *Escherichia coli*, with a focus on Shiga toxin-producing serotypes, global outbreaks (1982–2024) and food safety criteria. Gut Pathog 16:57. 10.1186/s13099-024-00641-9.

16. Ferreira C, Otani S, Aarestrup FM, Manaia CM. 2023. Quantitative PCR versus metagenomics for monitoring antibiotic resistance genes: balancing high sensitivity and broad coverage. FEMS Microbes 4:xtad008. 10.1093/femsmc/xtad008.

17. Bansal AK. 2005. Bioinformatics in microbial biotechnology–a mini review. Microb Cell Fact 4:19. 10.1186/1475-2859-4-19.

18. Kawanishi T, Shiraishi T, Okano Y, Sugawara K, Hashimoto M, Maejima K, Komatsu K, Kakizawa S, Yamaji Y, Hamamoto H, Oshima K, Namba S. 2011. New detection systems of bacteria using highly selective media designed by SMART: selective medium-design algorithm restricted by two constraints. PLoS One 6:e16512. 10.1371/journal.pone.0016512.

19. Mallona I, Soneson C, Carrillo B, Lütge A, Incicau D, Gerber R, Sonrel A, Robinson MD. 2025. Building a continuous benchmarking ecosystem in bioinformatics. PLoS Comput Biol 21:e1013658. 10.1371/journal.pcbi.1013658.

20. Meyer F, Robertson G, Deng ZL, Koslicki D, Gurevich A, McHardy AC. 2025. CAMI Bench-marking Portal: online evaluation and ranking of metagenomic software. Nucleic Acids Res 53:W102–W109. 10.1093/nar/gkaf369.

21. Page AJ, Alikhan NF, Carleton HA, Seemann T, Keane JA, Katz LS. 2017. Comparison of classical multi-locus sequence typing software for next-generation sequencing data. Microb Genom 3:e000124. 10.1099/mgen.0.000124.

22. Tiwari SK, Ponsero AJ, Haynes S, Tálas J, Grimes KP, Telatin A. 2026. A Systematic Bench-mark of Antibiotic Resistance Gene Detection Tools for Shotgun Metagenomic Datasets. bioRxiv. 10.64898/2026.02.04.703716.

23. McIntyre ABR, Ounit R, Afshinnekoo E, Prill RJ, Hénaff E, Alexander N, Minot SS, Danko D, Foox J, Ahsanuddin S, Tighe S, Hasan NA, Subramanian P, Moffat K, Levy S, Lonardi S, Greenfield N, Colwell RR, Rosen GL, Mason CE. 2017. Comprehensive benchmarking and ensemble approaches for metagenomic classifiers. Genome Biol 18:182. 10.1186/s13059-017-1299-7.

24. Yorki S, Shea T, Cuomo CA, Walker BJ, LaRocque RC, Manson AL, Earl AM, Worby CJ. 2023. Comparison of long- and short-read metagenomic assembly for low-abundance species and resistance genes. Brief Bioinform 24:bbad050. 10.1093/bib/bbad050.

25. Arabi H, Pakzad I, Nasrollahi A, Hosainzadegan H, Azizi Jalilian F, Taherikalani M, Samadi N, Monadi Sefidan A. 2015. Sulfonamide Resistance Genes (sul) M in Extended Spectrum Beta Lactamase (ESBL) and Non-ESBL Producing *Escherichia coli* Isolated From Iranian Hospitals. Jundishapur J Microbiol 8:e19961. 10.5812/jjm.19961v2.

26. Bourdonnais E, Colcanap D, Le Bris C, Brauge T, Midelet G. 2022. Occurrence of Indicator Genes of Antimicrobial Resistance Contamination in the English Channel and North Sea Sectors and Interactions With Environmental Variables. Front Microbiol 13:883081. 10.3389/fmicb.2022.883081.

27. Wang X, Yu D, Chui L, Zhou T, Feng Y, Cao Y, Zhi S. 2024. A Comprehensive Review on Shiga Toxin Subtypes and Their Niche-Related Distribution Characteristics in Shiga-Toxin-Producing E. coli and Other Bacterial Hosts. Microorganisms 12:687. 10.3390/microorganisms12040687.

28. Wirth T, Falush D, Lan R, Colles F, Mensa P, Wieler LH, Karch H, Reeves PR, Maiden MCJ, Ochman H, Achtman M. 2006. Sex and virulence in *Escherichia coli*: an evolutionary perspective. Mol Microbiol 60:1136–1151. 10.1111/j.1365-2958.2006.05172.x.

29. Gourlé H, Karlsson-Lindsjö O, Hayer J, Bongcam-Rudloff E. 2019. Simulating Illumina metagenomic data with InSilicoSeq. Bioinformatics 35:521–522. 10.1093/bioinformatics/bty630.

30. Gordeev V, Hölzer M, Desirò D, Goraichuk IV, Knyazev S, Solo-Gabriele H, Skums P, Karthikeyan S, Evans A, Agrawal S, Lucaci AG, Mason CE, Su JM, Gibas C, Nagarajan N, Peres da Silva R, Drabcinski N, Munteanu V, Zhan L, Rubin J, Wu NC, Trister A, Ciorba D, Bostan V, Lobiuc A, Covasa M, Ophoff RA, Zelikovsky A, Dimian M, Mangul S. 2025. Leveraging wastewater sequencing to strengthen global public health surveillance. BMC Glob Public Health 3:23. 10.1186/s44263-025-00138-w.

31. Al-Mustapha AI, Tiwari A, Laukkanen-Ninios R, Lehto KM, Oikarinen S, Lipponen A, Pitkänen T, Heikinheimo A, WastPan Study Group. 2025. Wastewater based genomic surveillance key to population level monitoring of AmpC/ESBL producing *Escherichia coli*. Sci Rep 15:7400. 10.1038/s41598-025-91516-9.

32. Merda D, Vila-Nova M, Bonis M, Boutigny AL, Brauge T, Cavaiuolo M, Cunty A, Regnier A, Sayeb M, Vingadassalon N, Yvon C, Chesnais V. 2024. Unraveling the impact of genome assembly on bacterial typing: a one health perspective. BMC Genom 25:1059. 10.1186/s12864-024-10982-z.

33. Jolley KA, Bray JE, Maiden MCJ. 2018. Open-access bacterial population genomics: BIGSdb software, the PubMLST.org website and their applications. Wellcome Open Res 3:124. 10.12688/wellcomeopenres.14826.1.

34. Ingle DJ, Valcanis M, Kuzevski A, Tauschek M, Inouye M, Stinear T, Levine MM, Robins-Browne RM, Holt KE. 2016. *In silico* serotyping of *E. coli* from short read data identifies limited novel O-loci but extensive diversity of O:H serotype combinations within and between pathogenic lineages. Microb Genom 2:e000064. 10.1099/mgen.0.000064.

35. Alcock BP, Huynh W, Chalil R, Smith KW, Raphenya AR, Wlodarski MA, Edalatmand A, Petkau A, Syed SA, Tsang KK, Baker SJC, Dave M, McCarthy MC, Mukiri KM, Nasir JA, Golbon B, Imtiaz H, Jiang X, Kaur K, Kwong M, Liang ZC, Niu KC, Shan P, Yang JYJ, Gray KL, Hoad GR, Jia B, Bhando T, Carfrae LA, Farha MA, French S, Gordzevich R, Rachwalski K, Tu MM, Bordeleau E, Dooley D, Griffiths E, Zubyk HL, Brown ED, Maguire F, Beiko RG, Hsiao WWL, Brinkman FSL, Van Domselaar G, McArthur AG. 2023. CARD 2023: expanded curation, support for machine learning, and resistome prediction at the Comprehensive Antibiotic Resistance Database. Nucleic Acids Res 51:D690–D699. 10.1093/nar/gkac920.

36. Zhou S, Liu B, Zheng D, Chen L, Yang J. 2025. VFDB 2025: an integrated resource for exploring anti-virulence compounds. Nucleic Acids Res 53:D871–D877. 10.1093/nar/gkae968.

37. Clausen PTLC, Hallgren MB, Overballe-Petersen S, Marcelino VR, Hasman H, Aarestrup FM. 2025. Assembly-free typing of Nanopore and Illumina data through proximity scoring with KMA. NAR Genom Bioinform 7:lqaf116. 10.1093/nargab/lqaf116.

38. Li D, Liu CM, Luo R, Sadakane K, Lam TW. 2015. MEGAHIT: an ultra-fast single-node solution for large and complex metagenomics assembly via succinct de Bruijn graph. Bioinformatics 31:1674–1676. 10.1093/bioinformatics/btv033.

39. Kang DD, Li F, Kirton E, Thomas A, Egan R, An H, Wang Z. 2019. MetaBAT 2: an adaptive binning algorithm for robust and efficient genome reconstruction from metagenome assemblies. PeerJ 7:e7359. 10.7717/peerj.7359.

40. Sieber CMK, Probst AJ, Sharrar A, Thomas BC, Hess M, Tringe SG, Banfield JF. 2018. Recovery of genomes from metagenomes via a dereplication, aggregation and scoring strategy. Nat Microbiol 3:836–843. 10.1038/s41564-018-0171-1.

41. Weber LM, Saelens W, Cannoodt R, Soneson C, Hapfelmeier A, Gardner PP, Boulesteix AL, Saeys Y, Robinson MD. 2019. Essential guidelines for computational method benchmarking. Genome Biol 20:125. 10.1186/s13059-019-1738-8.

42. Inouye M, Dashnow H, Raven LA, Schultz MB, Pope BJ, Tomita T, Zobel J, Holt KE. 2014. SRST2: Rapid genomic surveillance for public health and hospital microbiology labs. Genome Med 6:90. 10.1186/s13073-014-0090-6.

43. Feldgarden M, Brover V, Gonzalez-Escalona N, Frye JG, Haendiges J, Haft DH, Hoffmann M, Pettengill JB, Prasad AB, Tillman GE, Tyson GH, Klimke W. 2021. AMRFinderPlus and the Reference Gene Catalog facilitate examination of the genomic links among antimicrobial resistance, stress response, and virulence. Sci Rep 11:12728. 10.1038/s41598-021-91456-0.

44. Seemann T. 2015. Abricate. https://github.com/tseemann/abricate.

45. Joensen KG, Scheutz F, Lund O, Hasman H, Kaas RS, Nielsen EM, Aarestrup FM. 2014. Real-time whole-genome sequencing for routine typing, surveillance, and outbreak detection of verotoxigenic Escherichia coli. J Clin Microbiol 52:1501–1510. 10.1128/JCM.03617-13.

46. Seemann T. 2015. mlst. https://github.com/tseemann/mlst.

47. Larsen MV, Cosentino S, Rasmussen S, Friis C, Hasman H, Marvig RL, Jelsbak L, Sicheritz-Pontén T, Ussery DW, Aarestrup FM, Lund O. 2012. Multilocus sequence typing of total-genome-sequenced bacteria. J Clin Microbiol 50:1355–1361. 10.1128/JCM.06094-11.

48. Bessonov K, Laing C, Robertson J, Yong I, Ziebell K, Gannon VPJ, Nichani A, Arya G, Nash JHE, Christianson S. 2021. ECTyper: *in silico Escherichia coli* serotype and species prediction from raw and assembled whole-genome sequence data. Microb Genom 7:000728. 10.1099/mgen.0.000728.

49. Joensen KG, Tetzschner AMM, Iguchi A, Aarestrup FM, Scheutz F. 2015. Rapid and Easy In Silico Serotyping of Escherichia coli Isolates by Use of Whole-Genome Sequencing Data. J Clin Microbiol 53:2410–2426. 10.1128/JCM.00008-15.

50. Wick R. 2019. Badread: simulation of error-prone long reads. J Open Source Softw 4:1316.10.21105/joss.01316.

51. Page AJ, Keane JA. 2018. Rapid multi-locus sequence typing direct from uncorrected long reads using Krocus. PeerJ 6:e5233. 10.7717/peerj.5233.

52. Quince C, Walker AW, Simpson JT, Loman NJ, Segata N. 2017. Shotgun metagenomics, from sampling to analysis. Nat Biotechnol 35:833–844. 10.1038/nbt.3935.

53. Gwinn M, MacCannell D, Armstrong GL. 2019. Next-Generation Sequencing of Infectious Pathogens. JAMA 321:893–894. 10.1001/jama.2018.21669.

54. Lakin SM, Dean C, Noyes NR, Dettenwanger A, Ross AS, Doster E, Rovira P, Abdo Z, Jones KL, Ruiz J, Belk KE, Morley PS, Boucher C. 2017. MEGARes: an antimicrobial resistance database for high throughput sequencing. Nucleic Acids Res 45:D574–D580. 10.1093/nar/gkw1009.

55. Perreten V, Boerlin P. 2003. A New Sulfonamide Resistance Gene (sul3) in *Escherichia coli* Is Widespread in the Pig Population of Switzerland. Antimicrob Agents Chemother 47:1169–1172. 10.1128/aac.47.3.1169-1172.2003.

56. Wu S, Dalsgaard A, Hammerum AM, Porsbo LJ, Jensen LB. 2010. Prevalence and characterization of plasmids carrying sulfonamide resistance genes among *Escherichia coli* from pigs, pig carcasses and human. Acta Vet Scand 52:47. 10.1186/1751-0147-52-47.

57. Blanco M, Schumacher S, Tasara T, Zweifel C, Blanco JE, Dahbi G, Blanco J, Stephan R. 2005. Serotypes, intimin variants and other virulence factors of eae positive Escherichia coli strains isolated from healthy cattle in Switzerland. Identification of a new intimin variant gene (eae-⌘2). BMC Microbiol 5:23. 10.1186/1471-2180-5-23.

58. McDaniel TK, Jarvis KG, Donnenberg MS, Kaper JB. 1995. A genetic locus of enterocyte effacement conserved among diverse enterobacterial pathogens. Proc Natl Acad Sci U S A 92:1664–1668. 10.1073/pnas.92.5.1664.

59. Hacker J, Carniel E. 2001. Ecological fitness, genomic islands and bacterial pathogenicity. A Darwinian view of the evolution of microbes. EMBO Rep 2:376–381. 10.1093/embo-reports/kve097.

60. Dobrindt U, Hochhut B, Hentschel U, Hacker J. 2004. Genomic islands in pathogenic and environmental microorganisms. Nat Rev Microbiol 2:414–424. 10.1038/nrmicro884.

61. Scholz M, Ward DV, Pasolli E, Tolio T, Zolfo M, Asnicar F, Truong DT, Tett A, Morrow AL, Segata N. 2016. Strain-level microbial epidemiology and population genomics from shotgun metagenomics. Nat Methods 13:435–438. 10.1038/nmeth.3802.

62. Anyansi C, Straub TJ, Manson AL, Earl AM, Abeel T. 2020. Computational methods for strain-level microbial detection in colony and metagenome sequencing data. Front Microbiol 11:1925. 10.3389/fmicb.2020.01925.

