## Supplementary Material for "The impact of strain heterogeneity on genomic profiling accuracy in mixed-strain *Escherichia coli* metagenomes"

### Supplementary Materials

This supplementary section provides supporting information for the analyses presented in the main text. Table S1 summarises the genomes used to construct the simulated metagenomic datasets, including associated antimicrobial resistance genes, virulence genes, sequence types, and serotypes. Supplementary Figures S1 and S2 illustrate the strain and serotype abundance distributions used to generate datasets with varying levels of community complexity. Supplementary Table S2 provides a summary of MLST prediction outputs across the evaluated tools. Additional tables (Tables S3–S6) report detailed serotyping performance metrics across the simulated abundance gradients, including O-antigen recovery, H-antigen recovery, overall antigen recovery, and complete serotype recovery.

**Table S1:** Metadata on whole genomes used for simulation of metagenomes. A total of 49 genomes were included across the ARG, virulence gene (VG), serotype, and MLST datasets.

| Dataset | Genome | Genes of Interest | ST | Serotype |
|---|---|---|---|---|
| **ARG** | GCF_002002545 | blaTEM, sul2 | | |
| **ARG** | GCF_002001905 | blaTEM, sul1, sul2, tetA | | |
| **ARG** | GCF_002002175 | blaTEM, sul2 | | |
| **ARG** | GCF_002001715 | blaTEM, sul1, sul2, tetA | | |
| **ARG** | GCF_002002285 | blaTEM, sul2, tetA | | |
| **ARG** | GCF_002002075 | blaTEM, sul2 | | |
| **ARG** | GCF_002002235 | blaTEM, tetA | | |
| **ARG** | GCF_002001885 | blaTEM, sul2 | | |
| **ARG** | GCF_002002465 | blaTEM | | |
| **ARG** | GCF_002002315 | blaTEM | | |
| **ARG** | GCF_002001595 | blaTEM | | |
| **ARG** | GCF_002002165 | blaTEM, sul2 | | |
| **ARG** | GCF_002002125 | blaTEM, sul2, tetB | | |
| **ARG** | GCF_002001745 | blaTEM | | |
| **ARG** | GCF_002001665 | blaTEM, sul3, tetB | | |
| **ARG** | GCF_002002805 | blaTEM, sul2, tetB | | |
| **ARG** | GCF_002001525 | blaTEM | | |
| **ARG** | GCF_002001945 | blaTEM, tetA | | |
| **ARG** | GCF_002001585 | blaTEM, sul1, sul2, tetB | | |
| **ARG** | GCF_002001545 | blaTEM, sul1, sul2, tetA | | |
| **ARG** | GCF_002002225 | blaTEM, sul2 | | |
| **ARG** | GCF_002001705 | blaTEM | | |
| **ARG** | GCF_002001975 | blaTEM | | |
| **ARG** | GCF_002002105 | blaTEM, sul2, tetD | | |
| **ARG** | GCF_002001535 | blaTEM, tetB | | |
| **ARG** | GCF_002001785 | blaTEM, sul1, sul2, tetA | | |
| **ARG** | GCF_002002255 | blaTEM | | |
| **ARG** | GCF_002001855 | blaTEM, sul2, tetA | | |
| **ARG** | GCF_002002585 | blaTEM, sul1, sul2, tetA | | |

*Continued on next page*

*Table S1 continued from previous page*

| Dataset | Genome | Genes of Interest | ST | Serotype |
|---|---|---|---|---|
| **ARG** | GCF_002001915 | blaTEM, sul1, sul2, tetA | | |
| **ARG** | GCF_002001845 | blaTEM, sul3, tetA | | |
| **VG** | GCF_000520035 | stx2a, eae | | |
| **VG** | GCF_000010765 | stx1, stx2a, eae | | |
| **VG, ST** | GCF_000091005 | stx1, eae | 21 | O26:H11 |
| **VG, ST** | GCF_000010745 | stx1, stx2a, eae | 17 | O103:H2 |
| **VG, ST** | GCF_000008865 | stx1, stx2a, eae | 11 | O157:H7 |
| **VG, ST** | ecoli_k1_904 | k1 | 95 | O1:K1 |
| **VG, ST** | ecoli_k1_923. | k1 | 95 | O1:K1 |
| **VG, ST** | ecoli_k1_939 | k1 | 95 | O1:K1 |
| **ST** | ECOR10 | | 43 | O6:H10 |
| **ST** | ECOR11 | | 10 | O6:H10 |
| **ST** | ECOR12 | | 10 | O7:H32 |
| **ST** | ECOR26 | | 53 | O104:H21 |
| **ST** | ECOR36 | | 59 | O79:H25 |
| **ST** | ECOR54 | | 73 | O25:H1 |
| **ST** | ECOR55 | | 73 | O25:H1 |
| **ST** | ECOR56 | | 73 | O6:H1 |
| **ST** | ECOR59 | | 12 | O4:H40 |
| **ST** | ECOR72 | | 23 | O144:H8 |
| **ST** | ecoli_1359 | | 636 | O26 |
| **ST** | ecoli_1354 | | 833 | O26 |
| **ST** | ecoli_1362 | | 512 | O26 |
| **ST** | ecoli_1363 | | 77 | O83 |
| **ST** | ecoli_1356 | | 563 | O26 |

**Table S2:** Summary performance of MLST prediction tools across ten simulated metagenomes. Accuracy was calculated as the proportion of samples for which the predicted sequence type matched the dominant PCR-validated sequence type. All tools produced only unassigned sequence type calls, resulting in zero successful sequence type assignments.

| | **Assembly-based** | | **Read-based** | |
|---|---|---|---|---|
| Metric | **mlst** | **mlst-cge** | **metaMLST** | **SRST2** |
| Samples analysed | 10 | 10 | 10 | 10 |
| ST assignments | 0 | 0 | 0 | 0 |
| Unassigned ST calls | 10 | 10 | 10 | 10 |
| Correct ST assignments | 0 | 0 | 0 | 0 |
| ST assignment accuracy | 0% | 0% | 0% | 0% |

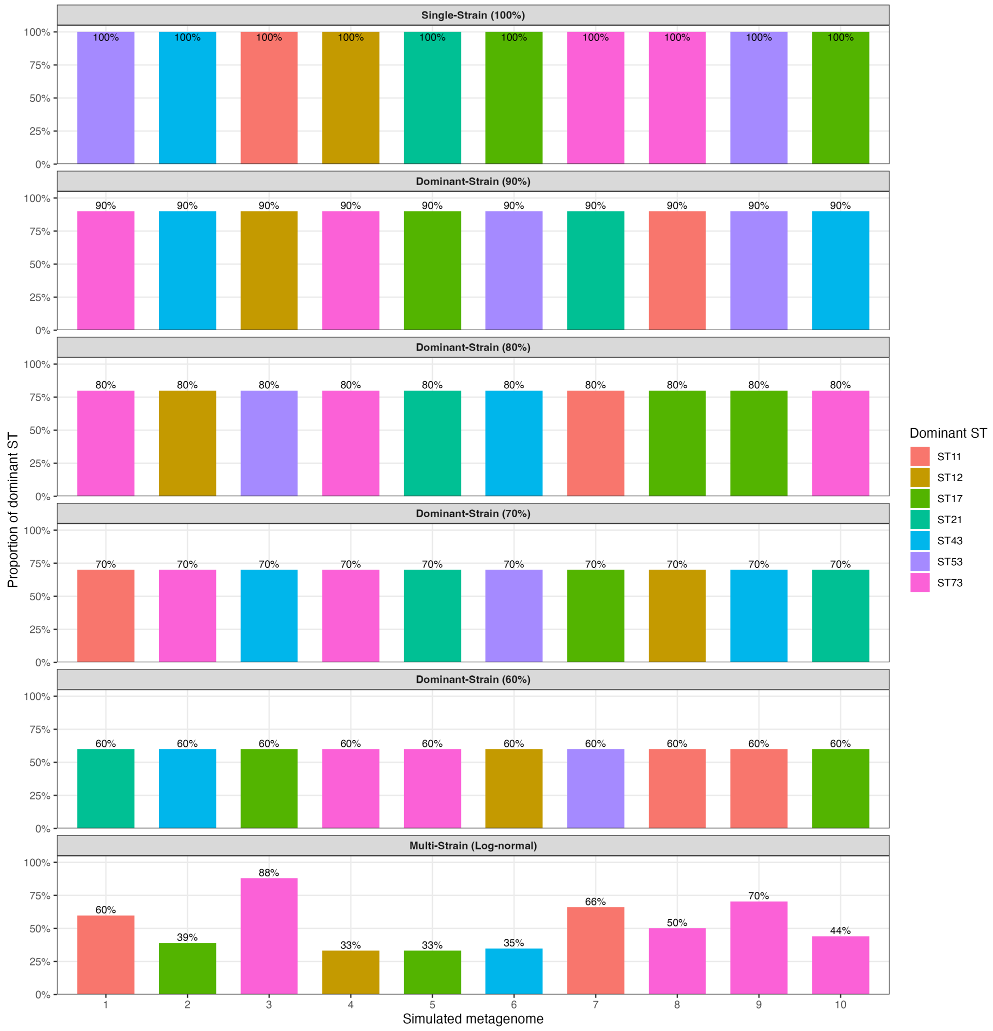


**Figure S1:** Dominant sequence type abundance across simulated metagenomic datasets. The single-strain dataset was generated with a single sequence type comprising 100% of reads. Dominant-strain datasets contained a dominant sequence type between 60% and 90% relative abundance in 10% intervals, while the multi-strain dataset was generated using log-normal abundance distributions. Bars indicate the proportion of the dominant sequence type within each simulated metagenome.

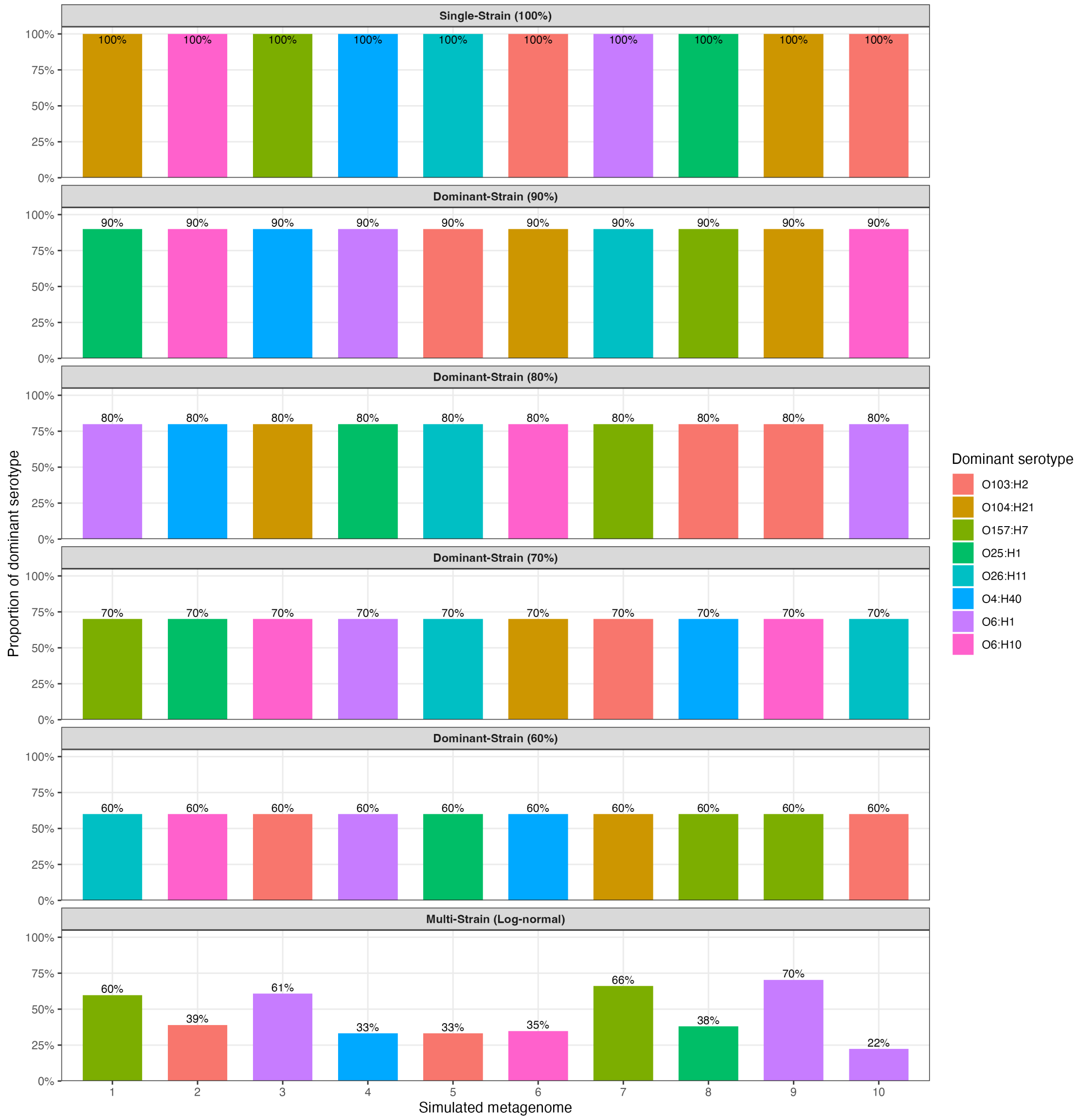


**Figure S2:** Dominant serotype abundance across simulated metagenomic datasets. The single-strain dataset was generated with a single serotype comprising 100% of reads. Dominant-strain datasets contained a dominant serotype between 60% and 90% relative abundance in 10% intervals, while the multi-strain dataset was generated using log-normal abundance distributions. Bars indicate the proportion of the dominant serotype within each simulated metagenome.

**Table S3:** Overall antigen recovery rates for serotype prediction tools across dataset compositions. Recovery rates were calculated as the proportion of expected O- and H-antigens successfully recovered across all datasets within each composition.

| Dataset | Assembly-based | | Read-based | |
|---|---|---|---|---|
| | ECTyper | SerotypeFinder | ECTyper | SRST2 |
| Single-strain (100%) | 90.0% | 95.0% | 35.0% | 95.0% |
| Dominant-strain (90%) | 85.0% | 90.0% | 30.0% | 90.0% |
| Dominant-strain (80%) | 70.0% | 80.0% | 30.0% | 75.0% |
| Dominant-strain (70%) | 80.0% | 85.0% | 15.0% | 60.0% |
| Dominant-strain (60%) | 70.0% | 85.0% | 20.0% | 55.0% |
| Multi-strain | 70.0% | 85.0% | 20.0% | 60.0% |

**Table S4:** O-antigen recovery rates across dataset compositions. Recovery was calculated as the proportion of samples for which the expected O-antigen was recovered.

| Dataset | Assembly-based | | Read-based | |
|---|---|---|---|---|
| | ECTyper | SerotypeFinder | ECTyper | SRST2 |
| Single-strain (100%) | 80.0% | 100.0% | 30.0% | 100.0% |
| Dominant-strain (90%) | 90.0% | 90.0% | 30.0% | 90.0% |
| Dominant-strain (80%) | 80.0% | 80.0% | 40.0% | 90.0% |
| Dominant-strain (70%) | 70.0% | 80.0% | 20.0% | 70.0% |
| Dominant-strain (60%) | 80.0% | 80.0% | 20.0% | 50.0% |
| Multi-strain | 80.0% | 80.0% | 40.0% | 60.0% |

**Table S5:** H-antigen recovery rates across dataset compositions. Recovery was calculated as the proportion of samples for which the expected H-antigen was recovered.

| Dataset | Assembly-based | | Read-based | |
|---|---|---|---|---|
| | ECTyper | SerotypeFinder | ECTyper | SRST2 |
| Single-strain (100%) | 100.0% | 90.0% | 40.0% | 90.0% |
| Dominant-strain (90%) | 80.0% | 90.0% | 30.0% | 90.0% |
| Dominant-strain (80%) | 60.0% | 80.0% | 20.0% | 60.0% |
| Dominant-strain (70%) | 90.0% | 90.0% | 10.0% | 50.0% |
| Dominant-strain (60%) | 60.0% | 90.0% | 20.0% | 60.0% |
| Multi-strain | 60.0% | 90.0% | 0.0% | 60.0% |

**Table S6:** Complete serotype recovery rates across dataset compositions. Recovery was calculated as the proportion of samples for which both the expected O- and H-antigens were recovered.

| Dataset | Assembly-based | | Read-based | |
|---|---|---|---|---|
| | **ECTyper** | **SerotypeFinder** | **ECTyper** | **SRST2** |
| Single-strain (100%) | 80.0% | 90.0% | 10.0% | 90.0% |
| Dominant-strain (90%) | 80.0% | 90.0% | 0.0% | 80.0% |
| Dominant-strain (80%) | 50.0% | 70.0% | 0.0% | 60.0% |
| Dominant-strain (70%) | 70.0% | 70.0% | 0.0% | 40.0% |
| Dominant-strain (60%) | 50.0% | 70.0% | 0.0% | 40.0% |
| Multi-strain | 50.0% | 80.0% | 0.0% | 30.0% |
